# Bacteria Contribute Exopolysaccharides to an Algal-Bacterial Joint Extracellular Matrix

**DOI:** 10.1101/2023.09.27.559704

**Authors:** Valeria Lipsman, Olesia Shlakhter, Jorge Rocha, Einat Segev

## Abstract

Marine ecosystems are influenced by phytoplankton aggregation, which affects processes like marine snow formation and harmful events such as marine mucilage outbreaks. Phytoplankton secrete exopolymers, creating an extracellular matrix (ECM) that promotes particle aggregation. This ECM attracts heterotrophic bacteria, providing a nutrient-rich and protective environment. In terrestrial environments, bacterial colonization near primary producers relies on attachment and the formation of multidimensional structures like biofilms. Bacteria were observed attaching and aggregating within algal-derived exopolymers, but it is unclear if bacteria produce an ECM that contributes to this colonization. This study, using *Emiliania huxleyi* algae and *Phaeobacter inhibens* bacteria in an environmentally relevant model system, reveals a shared algal-bacterial ECM scaffold that promotes algal-bacterial aggregation. Algal exudates play a pivotal role in promoting bacterial colonization, stimulating bacterial exopolysaccharide (EPS) production, and facilitating a joint ECM formation. A bacterial biosynthetic pathway responsible for producing a succinoglycan-like compound contributing to bacterial ECM formation is identified. Genes from this pathway show increased expression in algal-rich environments. These findings highlight the underestimated role of bacteria in aggregate-mediated processes in marine environments, offering insights into algal-bacterial interactions and ECM formation, with implications for understanding and managing disturbances like marine mucilage events.

## Introduction

Microbial aggregation drives ecologically important processes in the marine environment^1,2^. The natural phenomenon of marine snow is caused by the aggregation of phytoplankton into small particles, driving the sinking of organic matter in the water column^3–6^. Marine snow plays a central role in surface-to-bottom fluxes, acting as a shuttle of nutrients from the photic zone to the bottom of the ocean^5^. In Mediterranean seas, water column stratification during summer and high nutrient levels promote the merging of these particles into massive gelatinous sheets, flocs and clouds, known as marine mucilage^7–10^. The mucilage poses several environmental threats, such as preventing oxygen diffusion to lower parts of the water column and suffocating entire coral reefs^9,10^. However, the factors contributing to marine mucilage formation and the microorganisms involved remain areas of limited understanding.

Aggregation of microorganisms was shown to benefit the aggregating cells by increasing the resistance to harmful substances, such as antimicrobial compounds^11^ and enabling utilization of complex nutrients^12,13^. Furthermore, microbial cooperation frequently relies on maintaining close proximity, facilitated by cluster formation^12^, and can lead to the development of multicellular life cycles in both prokaryotic and eukaryotic microorganisms^14–16^. Another benefit of aggregation was demonstrated in populations of the unicellular alga *Chlamydomonas*, in which aggregation serves as a defense mechanism in response to predation^17,18^.

Phytoplankton-derived polymers, such as exopolysaccharides (EPS), promote aggregation^19–21^. Transparent exopolymer particles (TEP) are acidic polysaccharides produced by phytoplankton^22^. These particles are enriched in sulfate half-ester groups that confer high surface activity^23^. As a result, TEP can contribute to the formation of a complex and interconnected extracellular matrix (ECM). The adhesiveness of TEP was previously demonstrated to facilitate aggregation during phytoplankton blooms^24,25^ and in laboratory cultures^19^. Additionally, TEP can provide a surface for bacterial attachment and colonization^20,22,26–28^, creating rich microenvironments by supplying bacteria with nutrients and sanctuary^29,30^.

Bacteria were previously shown to promote aggregation in several algal species^6,31,32^, by physical attachment to the algal cells^6,32,33^ and by stimulating algal TEP production^31,32,34^. Interestingly, increased aggregation of phytoplankton upon bacterial colonization is generally attributed to changes in phytoplankton TEP production. The challenges in tracing the origin of TEP within mixed algal-bacterial populations have often led to overlooking bacteria as potential contributors to microbial TEP formation. This is of particular interest since marine bacteria were previously shown to produce TEP by release of capsular material and EPS^35,36^. However, it is unclear whether bacteria actively produce TEP when colonizing phytoplankton, and the algal-bacterial aggregation process remains poorly understood.

In the terrestrial environment, there is comprehensive understanding of the centrality of EPS in the associations of heterotrophic bacteria with primary producers. Bacterial colonization of plant roots in the rhizosphere was demonstrated to promote the formation of multicellular structures such as bacterial biofilms^37–42^. Bacterial formation of plant-associated biofilms is a process which involves several distinct steps^43–46^. Initially, plant exudates secreted by the roots act as signals that promote chemotaxis by planktonic bacteria^47,48^. The early steps of colonization involve an initial, reversible attachment of bacteria to a surface, often stabilized by the secretion of specific bacterial proteins or exopolysaccharide adhesins^42,49–52^. Plant-derived polysaccharides were demonstrated to serve as an environmental signal that triggers the bacterial switch to form biofilms^53^. The formation of a mature biofilm is characterized by a distinct spatial organization and the production of an adhesive ECM^46^. EPS is a key ECM component in most biofilms, and is often central in the formation of multidimensional structures^54–56^. The ECM creates an enclosed environment important for establishing plant-bacteria interactions on and near roots^57,58^.

In marine phytoplankton-bacteria interactions, namely between microalgae and abundant Roseobacter bacteria, much knowledge has accumulated about the algal-bacterial metabolic exchange^59–61^. However, our understanding of algal-bacterial aggregation is partial at best. A handful of studies addressed the process of bacterial colonization and direct attachment to algal cells^62–65^. In these previous studies, the term biofilm is largely used to describe bacterial attachment and not an ECM-held multidimensional structure, as established in literature of terrestrial plant-bacteria systems. Many uncertainties remain regarding bacterial colonization and algal-bacterial aggregation, particularly when drawing insights from terrestrial systems. Despite exhibiting resemblances to other models of primary producer-bacterial associations, the ability of marine bacteria to form multidimensional ECM-based structures in response to algal signals is yet to be established. Additionally, the composition and source of the ECM responsible for agglomerating algal-bacterial aggregates remain unknown.

Here, we use an environmentally relevant marine model system to study algal-bacterial aggregation and their ECM. Our model system includes the algal strain *Emiliania huxleyi* and the bacterial species *Phaeobacter inhibens*. This algal-bacterial model system was previously studied by us^59,63,66–69^ and by others^70–74^, revealing many routes of interactions and their environmental importance. Bacteria of the *P. inhibens* species attach through their pole to both abiotic and biotic surfaces^62,67^, including direct attachment to *E. huxleyi* cells^59^. As established in terrestrial systems, attachment is the first step towards forming multidimensional ECM-held structures^40–42^. However, whether *P. inhibens* can produce ECM-held biofilms subsequent to attachment is unknown. In the current study, we aimed to understand the process of algal-bacterial aggregation by addressing several key questions: Do marine bacteria form an ECM when interacting with algae? Do algae promote bacterial organization into multidimensional ECM-agglomerated structures? Do algae and bacteria aggregate through a joint ECM? Is a joint ECM produced by algae or bacteria? Elucidating these central aspects of algal-bacterial aggregation can provide insight into natural phenomena such as marine snow, and enhance our understanding about instances of perturbed conditions that result in harmful aggregation, like marine mucilage. With the increased frequency of catastrophic marine mucilage events^8,75,76^, understanding the process of algal-bacterial aggregation is an important step towards developing applicable treatments for these environmental threats.

## Results

### Algal exudates promote bacterial surface attachment

Algal exudates attract bacteria through chemotaxis and promote aggregation around the algal cell^77,78^. It therefore appears plausible that algal exudates will promote the bacterial switch from a motile to a sessile lifestyle, characterized by surface attachment. To monitor bacterial surface attachment upon exposure to algal exudates, algal cultures of *E. huxleyi* (at mid exponential growth) were filtered to remove cells and collect the exudate-containing filtrate (termed spent medium). To study the effect of algal exudates on the gradual process of the bacterial transition from motility to surface attachment, we introduced the algal spent medium into *P. inhibens* bacterial cultures at early growth stages. Bacterial attachment was assessed during different growth phases (mid exponential, early stationary and stationary) using an attachment assay^79^.

Bacteria supplemented with algal exudates showed increased attachment compared to control cultures grown in artificial seawater (ASW) (Fig. 1A). Algal spent medium significantly increased bacterial attachment at all bacterial growth phases and across a range of exudate concentrations. Furthermore, during mid exponential and early stationary growth, bacterial attachment linearly correlated with the increase in algal exudate concentrations between 0-75%. These findings suggest that bacterial attachment in response to algal cues is concentration-dependent, similar to chemotaxis^80^.

**Figure 1.**
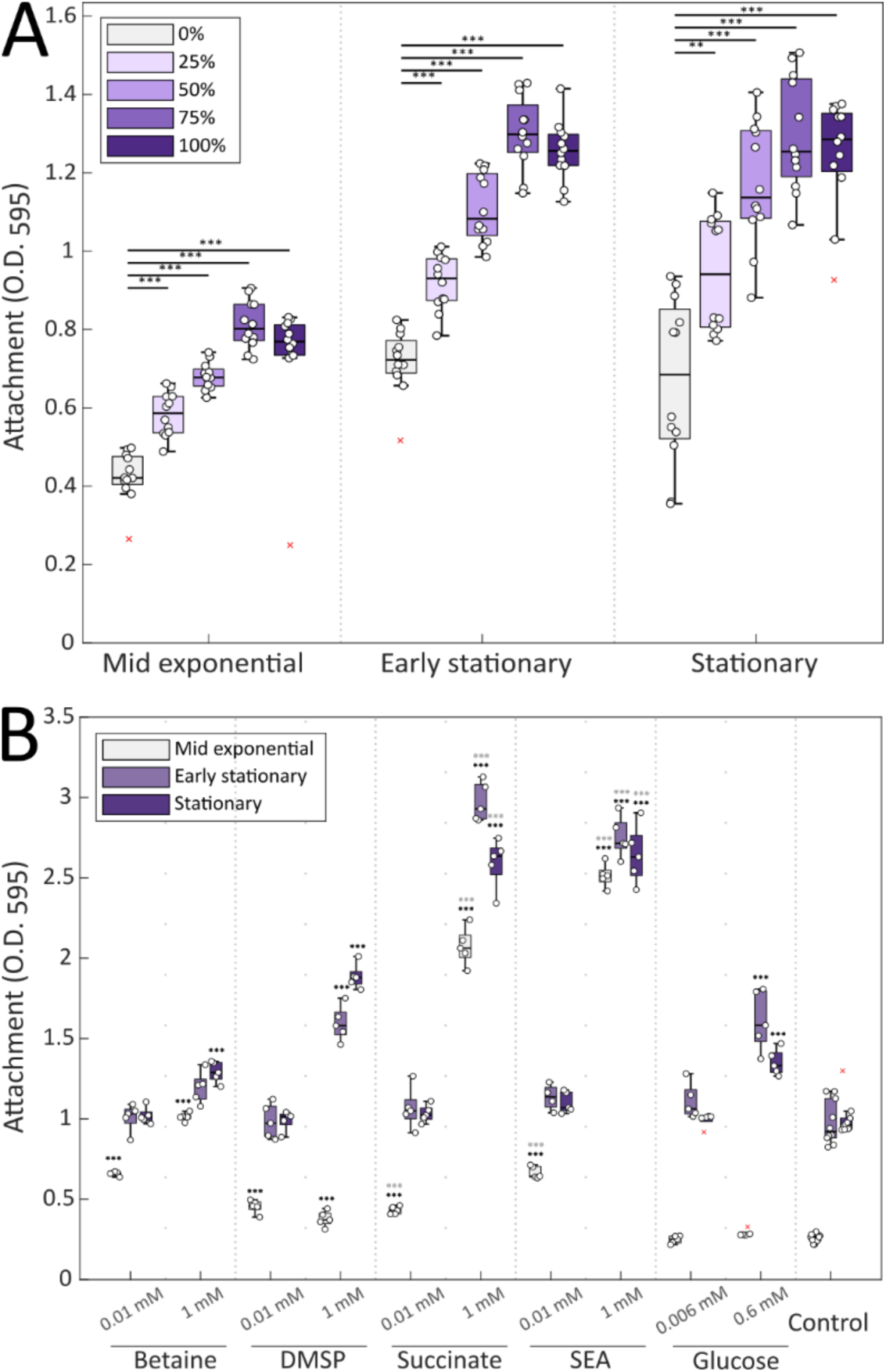
Algal exudates promote bacterial surface attachment. *P. inhibens* bacteria were grown with different supplements to evaluate the effect of algal exudates on surface attachment. Attachment was assessed measuring the absorbance of the Crystal Violet dye from bacterial cells that were strongly attached to 96-well plates^79^ (see Material and Methods) at O.D._595_. Bacteria at different growth phases (mid exponential, early stationary, and stationary) were assessed. Each sample was normalized to Crystal Violet extracted from the same medium and supplements but without bacteria. **(A)** Bacteria supplemented with different concentrations of algal spent medium (0-100% v/v). Spent medium was retrieved by filtering exponentially growing algal *E. huxleyi* cultures. Each box consists of n=12 wells. **(B)** Bacteria were supplemented with low (0.01 mM) or high (1 mM) concentrations of DMSP, betaine or succinate. Bacteria without added compounds were used as a control. Glucose was added as an additional control treatment at 0.006 mM or 0.6 mM, to account for the same amount of carbon present in other treatments. The synthetic exudates of algae (SEA) mix contained either 0.01 mM or 1 mM of each compound. Each box consists of n=5 wells. Box-plot elements are: center line - median; box limits - upper and lower quartiles; whiskers – min and max values, red x – outliers. Statistical significance was calculated using ANOVA with post-hoc Bonferroni test. Black asterisks mark treatments in which attachment was significantly higher than in ASW without any additional treatment, and at the same bacterial growth phase. Gray asterisks (in panel B) mark treatments of succinate or SEA that showed significantly higher attachment compared to treatments of glucose at the same bacterial growth phase. Two or three asterisk denote p-values lower than 0.01 or 0.001, respectively.

Algal spent medium contains metabolites that can facilitate bacterial growth^81–83^. To test whether improved bacterial attachment is the result of increased growth, we monitored bacterial growth until stationary phase in cultures supplemented with algal exudates versus ASW. in Our data showed that bacteria supplemented with spent medium exhibited earlier transition to logarithmic growth as previously reported^69^ (Fig. S1), however these cultures reached maximal cell numbers that are slightly lower than cultures grown in ASW. Thus, it appears that bacterial enhanced attachment response to algal exudates is dose-dependent, and the improved attachment does not stem from increased bacterial growth.

### Synthetic algal exudates promote bacterial attachment in a concentration-dependent manner

Algal exudates diffuse from the algal cell and create a gradient of exudates with high concentrations close to the cell and decreasing concentrations away from the algal cell ^77,84–86^. As spent medium represents the bulk average of algal exudates, it fails to recapitulate the elevated exudate concentrations that bacteria experience in proximity to the algal cell. To study the effect of high levels of algal exudates on bacterial attachment, we supplemented bacteria with selected algal exudates to allow precise manipulation of concentrations. Known algal exudates-dimethylsulfoniopropionate (DMSP), betaine and succinate-were previously monitored in laboratory algal cultures and in the environment^78,87,88^.

Importantly, these compounds were previously shown to influence the chemotactic behavior in various marine bacteria^78,86,89^. It was demonstrated that the concentrations of DMSP, for example, in bulk seawater are in the nanomolar range^90^, while algal intracellular concentration can be orders of magnitude higher^91^. To represent the chemical environment that bacteria may experience in different distances from the algal cell, bacterial cultures were supplemented with individual compounds in concentrations that represent proximity to the algal cell (1 mM) or distance from algae (0.01 mM).

Low concentrations (0.01 mM) of the individual compounds-DMSP, betaine or succinate-increased bacterial attachment when the treated bacterial cultures were at mid-exponential phase (Fig. 1B). High concentrations (1 mM) of these compounds further increased bacterial attachment to levels greater than the attachment values observed upon treatment with algal spent medium (Fig. 1A). DMSP, betaine and succinate can serve as an additional carbon source for bacteria^92–97^. To test whether increased attachment was achieved due to additional carbon availability, we supplemented bacterial cultures with glucose. The added glucose concentrations were chosen to reflect the carbon content of the succinate treatment, which showed the most pronounced improvement of attachment (Fig. 1B, 0.006 mM and 0.6 mM glucose, for low and high concentrations respectively). Results of these experiments demonstrate that the added glucose did not improve attachment as observed upon treatment with succinate. Thus, succinate is likely to act as a signal that promotes bacterial attachment rather than a carbon source. Moreover, the maximal cell numbers of bacteria that was reached upon treatment with succinate was lower than untreated bacteria (Fig. S2), further demonstrating that the improved attachment is not due to increased growth.

In the environment, bacteria encounter a mixture of molecules exuded by algae. Therefore, we assembled a synthetic mixture with the representative algal exudates (SEA-Synthetic Exudates of Algae) that contained either 1 mM or 0.01 mM of each individual compound (DMSP, betaine and succinate). Results demonstrate that 1 mM of the SEA mix significantly promotes attachment at mid exponential growth, beyond the levels observed for individual compounds. Thus, bacterial attachment is promoted by high concentrations of algal exudates, akin to the chemical environment in the proximity of the algal cell.

### Algal exudates promote bacterial organization in multidimensional structures

Bacterial surface attachment is the first necessary step in the formation of complex, multidimensional structures such as biofilms^43,46^. However, not all bacteria that possess attachment capabilities can organize in multidimensional structures, and thus remain attached in monolayers^54^. To determine whether conditions that promote attachment of *P. inhibens* bacteria, also induce multidimensional organization, we monitored the spatial organization of bacteria supplemented with 1 mM of the SEA mix. Specifically, bacteria were provided with glass slides as a surface for attachment, and the thickness of the bacterial structures that were formed was monitored using confocal microscopy (Fig. S3).

Bacteria supplemented with the SEA mix organized in significantly thicker structures compared to untreated bacteria (Fig. 2). At stationary phase, SEA-treated bacteria appeared to be organized in a continuous layer and exhibited significantly higher mean thickness up to 18 μm (Fig. 2A), while untreated bacteria attached in thin monolayers with dispersed bacterial patches on the glass surface (Fig. 2B). Therefore, it appears that algal exudates promote *P. inhibens* bacterial attachment and the subsequent organization in multidimensional structures.

**Figure 2.**
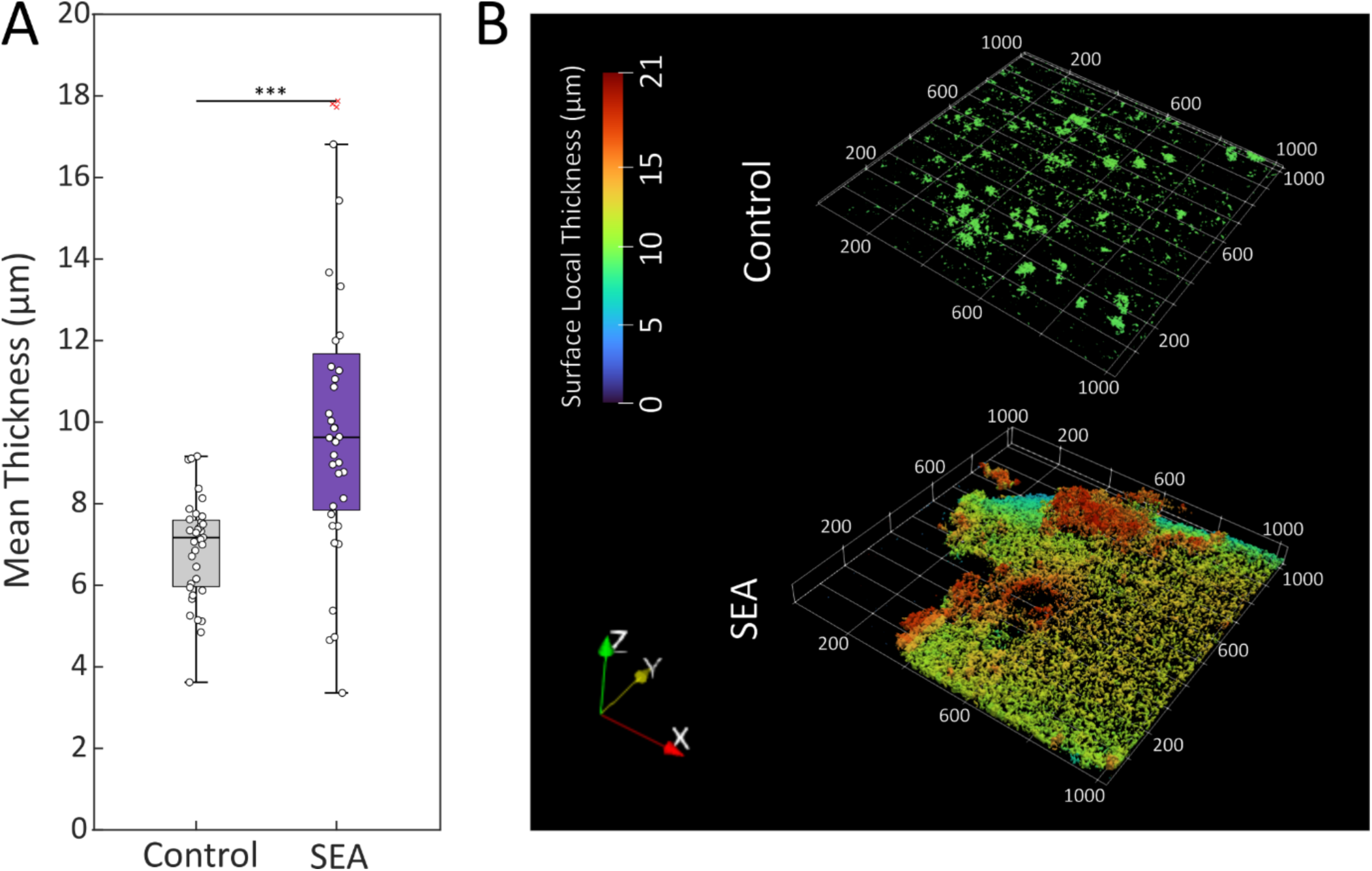
The SEA mix promotes bacterial organization in multidimensional structures. Multidimensional organization of *P. inhibens* bacteria was monitored using confocal imaging of Syto9-stained bacterial structures that formed on glass slides (see Materials and Methods). Bacterial cultures were supplemented with 1 mM of the SEA mix, or a corresponding volume of ASW as control, and grown to stationary phase. Multidimensional organization was quantified using BiofilmQ^194^ with z-stacks of the imaged slides. **(A)** Calculation of the mean thickness (µm) was conducted using 7 random points on 5 different slides. Box-plot elements are: center line - median; box limits - upper and lower quartiles; whiskers – min and max values, red x – outlier. Statistical significance was calculated using ANOVA with post-hoc Bonferroni test. Three asterisks mark statistically different results (p-value<0.001). **(B)** Example of multidimensional structures of bacteria at stationary phase that were supplemented with ASW as control (top) or SEA mix (bottom). The color bar corresponds to the surface local thickness (µm) used to calculate the mean thickness of each image.

### Algal exudates increase the production of bacterial EPS

Bacterial multidimensional structures are held together by an extracellular matrix (ECM)^54–56^. The bacterial ECM was largely studied in biofilms of terrestrial bacteria ^43–45^ and was shown to consist of macromolecules such as extracellular polysaccharides (EPSs), proteins and extracellular DNA, which serve as a scaffold^54–56^. In the marine environment, EPSs are of particular interest, as they are known to be produced by various phytoplankton species and play a key role in the formation of marine snow and the global carbon cycle^20,22,98–102^. To determine whether bacterial EPSs are involved in agglomerating the multidimensional structures of SEA-treated bacteria, we stained bacterial samples with the Alcian Blue dye that stains acidic polysaccharides, similar to EPS derived from algae^103^. Our results show that SEA-treated bacteria produce extracellular threads and sheet-like structures that are stained by the dye, while these structures are not observed in control cultures (Fig. S4A). Moreover, quantification of the extracellular carbohydrates extracted from SEA-treated bacteria showed significantly higher amounts of EPS compared to cultures of untreated bacteria (Fig. S4B). Therefore, it is likely that the polysaccharide-containing structures are part of the bacterial ECM that is involved in binding multidimensional structures, and its production is induced by algal exudates.

### Algal exudates promote expression of bacterial genes related to ECM production

SEA mix treatment promoted bacterial transition from motility to attachment and stimulated multidimensional organization through ECM production. To gain insights into these processes, we wished to examine the expression of the genetic modules that underlie these bacterial transitions. Therefore, we mined the literature and compiled a list of indicative genes that serve as markers for each process (Table S1). As motility markers, we chose *motA* and *motB* which have an established role in bacterial motility across various bacteria^104^. As a marker for initial surface attachment, we chose the gene *dltA1* which was shown to be involved in initial surface attachment of other bacteria, through the biosynthesis of d-alanyl-lipoteichoic acid (LTA)^105^. Although LTA is specific to Gram-positive bacteria, a few Gram-negative bacteria like *P. inhibens* carry this gene and its possible function in marine bacteria was previously discussed^106–108^. *P. inhibens* also carries several genes previously annotated as belonging to the biosynthetic pathway of succinoglycan, an EPS known to be produced by *Rhizobial*^109^ and *Agrobacterium*^110^ species. As markers for EPS production, we chose *exoB*, and two copies of genes annotated as *exoY*, found on the chromosome and the 262kb plasmid of *P. inhibens*. Both genes encode for proteins that are responsible for the priming of succinoglycan biosynthesis: *exoB* encodes for a protein which produces the activated nucleotide-sugar UDP-galactose^111^, which is later transferred by the protein encoded by *exoY* to a lipid carrier, marking the beginning of the primary sugar chain biosynthesis^109,112,113^.

We followed the expression of the marker genes over time under SEA mix treatment using qRT-PCR. Our data show that soon after supplementing the SEA mix (3 hours, during mid exponential growth), motility genes were downregulated (*motA* and *motB*), while genes potentially involved in surface attachment (*dltA1*) and biosynthesis of the activated nucleotide-sugar (*exoB*) were upregulated (Fig. 3). Later, at stationary phase (3 days), the gene *exoY_c_*, potentially involved in the production of succinoglycan, was upregulated, while the *dltA1* gene that is potentially involved in initial surface attachment was downregulated. Interestingly, only the chromosomal *exoY* was upregulated, and not the copy found on the plasmid, suggesting different regulation of these two gene copies. Thus, algal exudates appear to induce a genetic program in bacteria. The program initially triggers surface attachment and the early steps in production of specific EPS components and progresses over time to promote the production of these EPS components.

**Figure 3.**
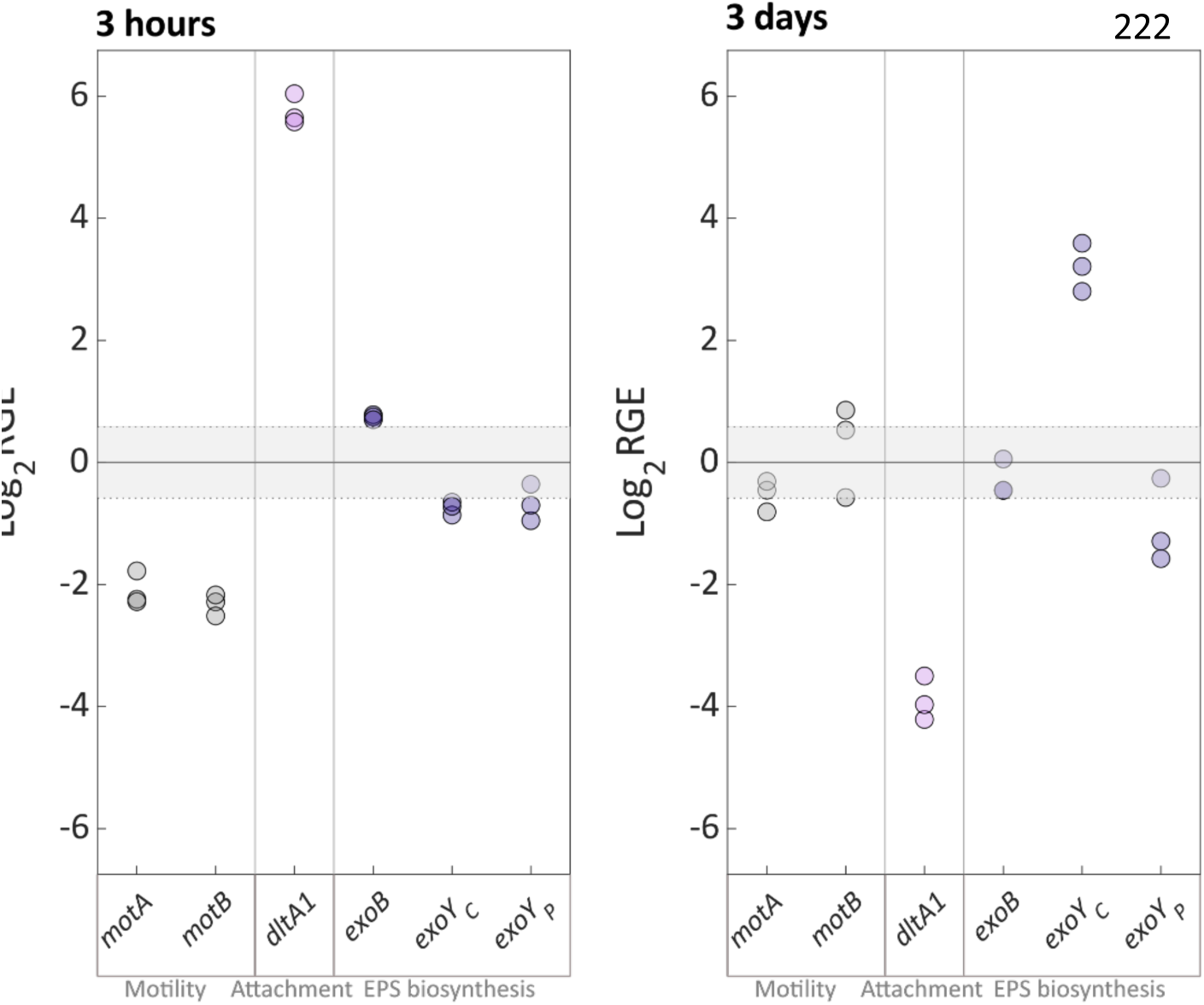
The SEA mix increases expression of bacterial genes potentially involved in attachment and ECM production. The relative gene expression (RGE) of genes involved in motility, attachment, and production of ECM-related EPS was measured 3 hours (mid exponential, left) or 3 days (stationary, right) after addition of the SEA mix. Gene expression was normalized to two housekeeping genes (*gyrA* and *recA*) and to untreated samples collected at the same time. RNA was collected from three biological replicates for both SEA-treated and untreated bacteria. Upregulation or downregulation was determined according to a threshold of ±1.5 RGE.

### Bacterial gene modules involved in EPS production are expressed in algal-bacterial co-cultures

To gain deeper insights into the production of bacterial EPS in the context of algal-bacterial interactions, we tracked the expression of bacterial genes potentially involved in EPS biosynthesis in algal-bacterial co-cultures. First, we curated the genetic modules that are potentially involved in the EPS biosynthetic pathway in *P. inhibens*. Functional annotation, using EggNOG-mapper v2^114,115^, revealed gene clusters around the two *exoY* genes (Fig. S5A), resembling other characterized wzx/wzy-dependent EPS modules^116^ (Table S2). Both *P. inhibens* EPS modules contained genes encoding proteins responsible for priming glycotransferases, primary sugar chain glycotransferases, repeating unit flippases, repeating unit polymerases and polysaccharide co-polymerases. The plasmid module also contained a gene that encodes for an outer membrane polysaccharide export protein, which was missing from the chromosomal module, but was found outside of the chromosomal gene cluster. Genes for other functional groups, such as activated sugar nucleotide biosynthesis, were found in other genomic locations similar to the architecture of other EPS modules^116^. We were unable to annotate genes encoding for functions related to modification of the sugar bases in the repeat units, such as acyl groups, found in other characterized EPSs^116^. However, many wzx/wzy-dependent EPSs, including succinoglycans from different bacterial species, show variability in acyl group composition and corresponding gene variability^116–118^. This variability could explain the lack of clear orthologs in the EPS modules of *P. inhibens*.

The reconstruction of the two EPS biosynthetic modules of *P. inhibens* bacteria now allowed us to track relevant gene expression in algal-bacterial co-cultures. Previously, our lab has generated transcriptomic data of *P. inhibens* bacteria in co-culture with *E. huxleyi* algae^66^. Mining these data revealed marked expression of bacterial genes encoding for EPS-biosynthesis related functions in co-cultures with algae. Specifically, during exponential growth, upregulation of the expression of *exoY*-like gene from the chromosomal module was evident in bacteria grown in co-cultures, compared to bacterial monocultures under similar growth phase (Fig. S5B). In contrary, the *exoY* gene from the plasmid module was not upregulated in co-cultures, suggesting different regulation from the chromosomal module. These gene expression patterns are similar to patterns observed in bacterial monocultures in response to SEA treatment (Fig. 3). Taken together, bacteria appear to activate the succinoglycan-like biosynthetic pathway in the presence of algae, and the chromosomal *exoY* might be a marker gene to follow the production of the succinoglycan-like EPS in *P. inhibens*.

### Algal exudates influence bacterial production of polysaccharides involved in attachment and ECM

Our observations suggest that bacterial EPSs play a role in attachment and ECM production of *P. inhibens* bacteria. To characterize the polysaccharides present in the extracellular milieu of SEA-treated bacteria, we used fluorophore-conjugated lectins to target and visualize specific sugar monomers. We first targeted the polar polysaccharide of *P. inhibens* bacteria which contains N-Acetyl-D-glucosamine (GlcNAc) residues and was previously shown to be involved in attachment between bacterial cells and abiotic surfaces^67,119–122^. Our observations revealed that untreated bacteria exhibit the characteristic polar staining as previously described^67^ (Fig. 4A, WGA-wheat germ agglutinin), while SEA-treated bacteria exhibit staining of the entire cell circumference (Fig. 4A and S6). These observations suggest that the production and cellular arrangement of the bacterial polysaccharide is influenced by algal exudates. Next, we wanted to characterize the *P. inhibens* extracellular EPS production, targeting the priming sugar that is added by the *exoY*-encoded proteins. Generally, the priming sugars in various wzx/wzy-dependent EPSs are either UDP-glucose or UDP-galactose, and their priming glycotransferases share the same functional annotation (Table S2). Aligning *P. inhibens exoY* genes to PANTHER HMM models^123^ revealed that the proteins encoded by the two *exoY* genes are from the family of colanic biosynthesis UDP-glucose lipid carrier transferases (PTHR30576, sub family SF10 for chromosomal *exoY*, and SF4 for plasmid *exoY*). Analyses of the protein sequences and the phylogenetic relationships (Fig. S7) of priming glycotranferases of characterized EPSs from various *Proteobacteria*, support that both proteins encoded by the *P. inhibens exoY* genes are responsible for priming galactose.

**Figure 4.**
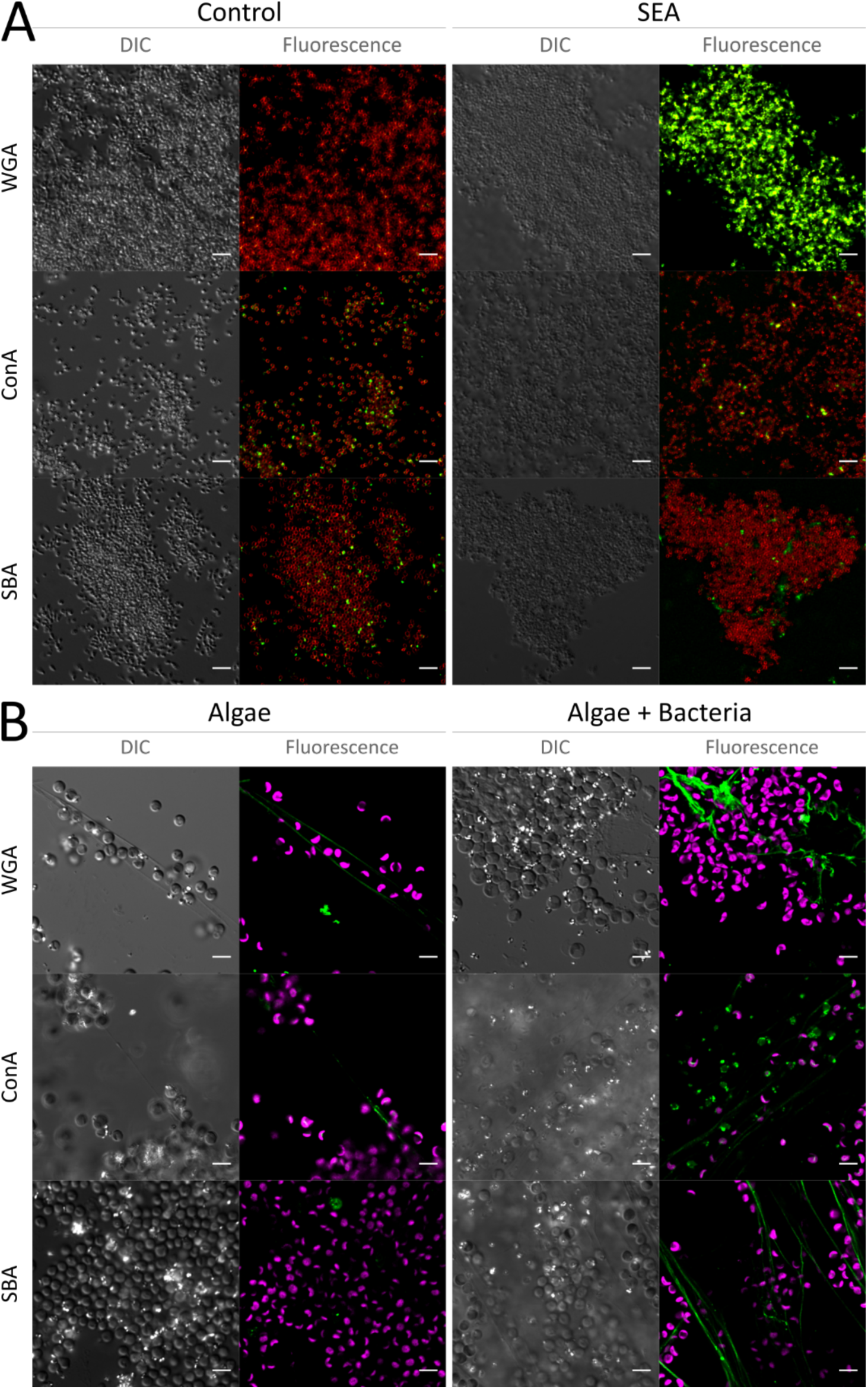
A bacterial EPS with succinoglycan characteristics is part of the ECM in bacterial cultures and algal-bacterial co-cultures. Confocal microscopy was used to study the sugar composition of the ECM of different cultures using a panel of FITC-conjugated lectins to detect sugar moieties (green in the fluorescence images). The lectins used were Wheat Germ Agglutinin (WGA) that binds to GlcNAc residues, Concanavalin A (ConA) that binds glucose residues, and Soybean Agglutinin (SBA) that binds galactose residues. **(A)** Bacterial *P. inhibens* pure cultures were grown to late stationary phase on glass slides and treated with SEA (right) or were untreated (left). Shown are DIC (Differential interference contrast) images for general cell abundance, the fluorescence of the bound FITC-conjugated lectins (green) and the membrane dye FM4-64 (red) that demarcates bacterial cells. **(B**) Algal *E. huxleyi* monocultures (left) and *E. huxleyi* - *P. inhibens* co-cultures were grown to late exponential phase and visualized. Shown are DIC images, the fluorescence of the bound FITC-conjugated lectins (green), and algal autofluorescence of chlorophyll *a* (pink). Scale bar corresponds to 5 µm.

Due to the high genetic resemblance between the *P. inhibens* EPS modules and previously characterized succinoglycan genetic modules, we focused on detecting extracellular EPS by targeting and visualizing the galactose and glucose residues^112,124^ using the fluorophore-conjugated lectins Soybean Agglutinin (SBA) and Concanavalin A (ConA), respectively. Importantly, SBA was previously shown to target the galactose residues in the EPS produced by *Rhizobium* species^125^, and EPS deficient mutants also displayed deficiency in SBA binding^126,127^. The ConA lectin showed similar binding to both SEA-treated and untreated bacteria, appearing as small extracellular foci (Fig.4A), suggesting that glucose is a common residue in substances secreted by the bacteria. However, SBA binding patterns appeared different between treated and untreated bacteria- in control cultures, binding occurred in small extracellular foci, while SEA-treated cultures exhibited a more dispersed signal that seemed to occupy the intercellular space (Fig. 4A and Fig. S8). These results suggest that *P. inhibens* bacteria produce an EPS with galactose residues - characteristic of succinoglycan - and this production is promoted by algal exudates.

### Algae and bacteria form a joint ECM with a unique EPS composition

Our results suggest that bacterial attachment and ECM production is enhanced when bacteria are exposed to algal exudates. How do these bacterial processes manifest when bacteria grow with live algae? Do they influence algal-bacterial co-cultures? On a macroscopic level, algal-bacterial co-cultures show visibly higher aggregation as the cultures age, while algal monocultures of the same age are more homogenous (Fig. S9). Microalgal cells, such as *E. huxleyi,* are known to exude transparent exopolymeric particles (TEP), characterized by Alcian Blue staining^22,128,129^. These polymers can self-assemble to form a dense, gelatinous and a continuous matrix, and can serve as a basis for bacterial attachment^22,28,130^. Our results show that in co-cultures, bacteria are often found entangled in exopolymeric substances (Fig. S10). Such exopolymers - in laboratory cultures and environmental samples - have been often considered to be of algal source^22,102,129,131,132^, originating from storage polysaccharides produced as a sink for excess carbon^98–100^. The role of bacteria in TEP distribution was previously investigated^101,133–135^, showing bacterial contribution to the marine TEP pool by various processes including release of EPS and capsular material^36^. Additionally, bacterial presence was previously shown to increase the production of TEP in cultures with phytoplankton^31,32,136,137^ Although bacteria can benefit from colonization of preexisting TEP^26,27,138^, it is not clear whether bacteria actively synthesize EPS in the algal environment. Since our data show that bacteria are capable of producing exopolymers in response to algal exudates, we investigated whether bacteria contribute to an algal-bacterial joint ECM. A composite ECM from both algal and bacterial sources might have a composition that can be attributed to both algal and bacterial origins. To characterize the sugar composition of the algal-bacterial ECM, the ECMs of algal monocultures and algal-bacterial co-cultures (Fig. 4B) were stained with fluorophore-conjugated lectins and visualized under the microscope.

Our results indicate that the GlcNAc moieties and glucose residues (recognized by the WGA and ConA lectins, respectively) are found in all ECMs-bacterial, algal and co-cultures (Fig. 4A and Fig. 4B). Therefore, the source of these compounds in the co-culture ECM currently cannot be determined. However, galactose residues were detected by the SBA lectin only in bacterial ECM (Fig. 4A) and algal-bacterial ECM (Fig. 4B). In the algal-bacterial ECM, the SBA lectin was bound to long extracellular filamentous structures that stretched tens of microns between cells. In contrast, in algal monocultures the SBA lectin showed rare binding to foci on algal cells. (Fig. 4B). These observations suggest that both algae and bacteria contribute to the EPS pool to form a joint ECM with a distinct sugar composition. Furthermore, these results support that when bacteria are in proximity to algal cells, they produce EPS with the characteristics of succinoglycan.

### The *exoY* gene plays a role in bacterial attachment

In *Sinorhizobium* and *Agrobacterium*, the *exoY* gene product is responsible for the initiation of the repeat subunit biosynthesis^109,112,113^. Importantly, mutation of *exoY* in *Sinorhizobium* completely abolishes the production of succinoglycan^139^, reducing the fraction of galactose residues in the EPS^140^. Thus, the upregulated expression of the chromosomal *exoY* upon SEA mix treatment (Fig. 3) and its high expression in co-cultures (Fig. S5B) point to a role of this gene in EPS production in *P. inhibens* bacteria. To assess the role of chromosomal *exoY* in EPS biosynthesis in *P. inhibens*, and its involvement in ECM production, we deleted the gene and generated the knockout mutant *ΔexoY*. We then tested the capability of *ΔexoY* bacteria to attach in response to SEA mix treatment. Deletion of *exoY* caused reduced attachment in both SEA-treated and untreated cells as compared to wild-type bacteria (WT) (Fig. 5A). These results suggest that the succinoglycan-like EPS plays a role in bacterial attachment but appears to function in concert with other attachment mechanisms, as *exoY* deletion reduced but did not abolish attachment. Importantly, growth curves of *ΔexoY* bacteria were similar to WT, both with SEA treatment and with no treatment (Fig. S11). These observations indicate that *exoY* deletion did not impair the ability of bacteria to respond to the compounds in the SEA mix.

**Figure 5.**
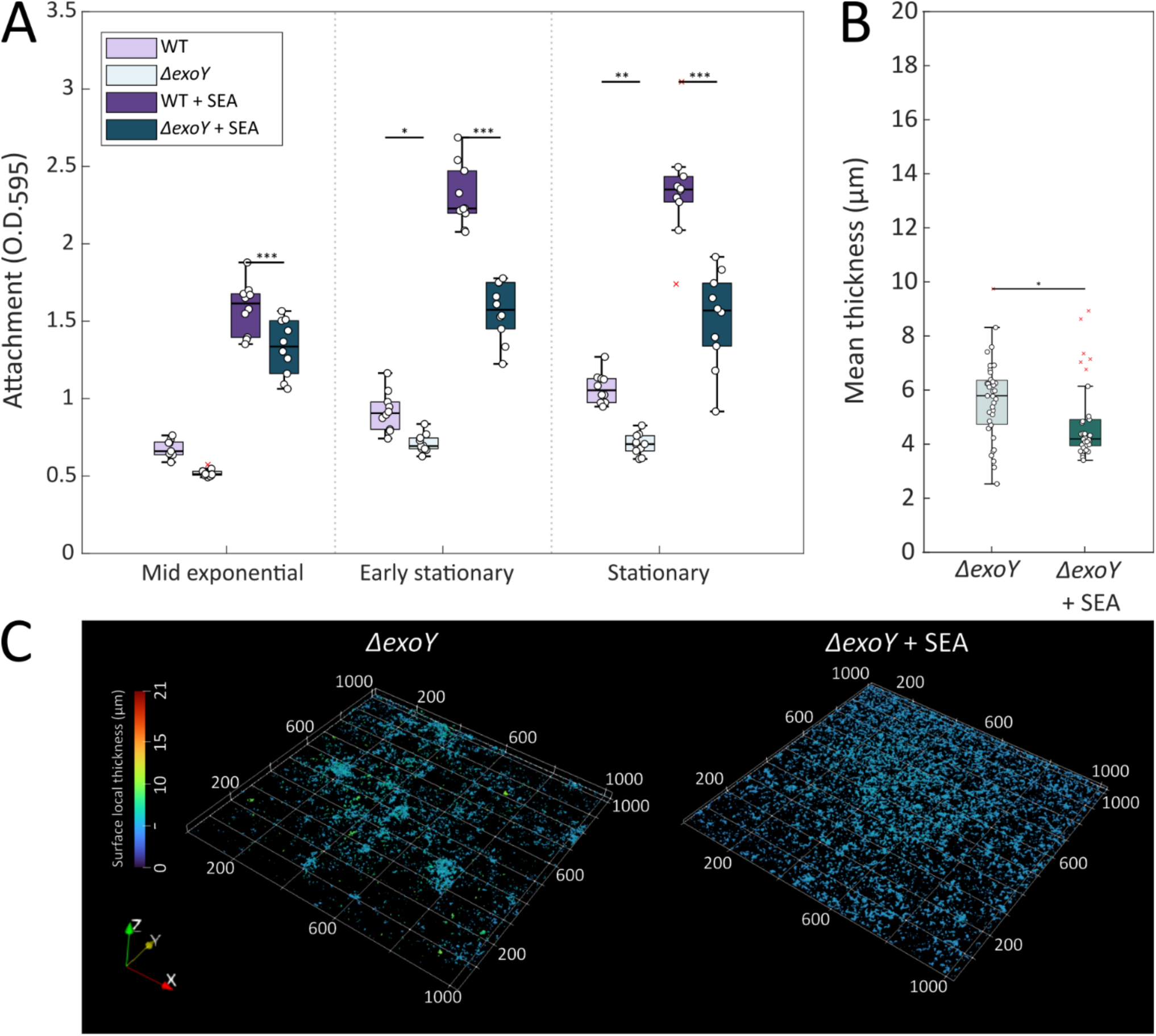
*P. inhibens* attachment capabilities and organization in multidimensional structures are reduced in *exoY* deletion mutants. **(A)** Attachment of *P. inhibens* WT (purple) or *ΔexoY* (green) bacteria was evaluated using an attachment assay (see Material and Methods). Bacteria were grown in ASW supplemented with SEA mix (1 mM of each molecule) or without added compounds as control. Attachment was assessed by extracting the Crystal Violet dye from bacterial cells strongly attached to 96-well plates and measuring absorbance at O.D._595_. Bacteria at different growth phases (mid exponential, early stationary, and stationary) were assessed. Each box consists of n=10 wells, normalized to Crystal Violet extracted from the same medium but without bacteria. Asterisks mark treatments in which significantly different attachment was observed between WT and *ΔexoY* bacteria at the same time point for the same treatment. **(B)** Multidimensional organization of *ΔexoY* bacteria was monitored using confocal imaging of Syto9-stained bacterial structures that formed on glass slides (see Materials and Methods). Cultures of *ΔexoY* bacteria were supplemented with 1 mM of the SEA mix or a corresponding volume of ASW as control and grown to stationary phase. Multidimensional organization was quantified using BiofilmQ^194^ with z-stacks of the imaged slides. Calculation of the mean thickness (µm) was conducted using random points (n>6) on 5 different slides. **(C)** Example of a multidimensional structure of *ΔexoY* bacteria at stationary phase that were supplemented with ASW as control (left) or SEA mix (right). The color bar corresponds to the surface local thickness (µm) used to calculate the mean thickness of each image. Box-plot elements are: center line - median; box limits - upper and lower quartiles; whiskers – min and max values, red x – outlier. Statistical significance was calculated using ANOVA with post-hoc Bonferroni test. One, two or three asterisks denote p-values lower than 0.05, 0.01 and 0.001, respectively.

### The *exoY* gene is important for bacterial multidimensional organization

Mutant *ΔexoY* bacteria showed significantly reduced surface attachment (Fig. 5A). Therefore, we further evaluated if *ΔexoY* mutants are perturbed in their ability to form multidimensional structures (Fig. 5B and C). Our results show that the multidimensional structures of the mutant bacteria were markedly disturbed under SEA treatment, appearing as sparse, small aggregates, with reduced local thickness (Fig. 5C). The SEA treatment slightly decreased the mean thickness of structures formed by mutant bacteria (Fig. 5B), while they both were significantly thinner than those of WT bacteria (Fig. 2A). These observations further highlight the centrality of the succinoglycan-like EPS as a scaffold material in the bacterial ECM.

### The *exoY* gene contributes to the bacterial ECM formation with a succinoglycan-like EPS

Next, we wished to validate that the perturbed attachment and spatial organization of the *ΔexoY* mutant bacteria is indeed due to the lack of extracellular EPS. Therefore, we monitored succinoglycan residues in the bacterial ECM of *ΔexoY* mutants using the fluorescently labeled ConA and SBA lectins (Fig. 6A). As can be seen in Fig. 6A, no sugar residues were detected in the extracellular environment of *ΔexoY* bacteria. Both glucose and galactose (detected by the ConA and SBA lectins, respectively) were not detected in the ECM of SEA-treated and untreated cultures. Staining of GlcNAc moieties in *ΔexoY* cultures showed similar patterns to WT staining, both in SEA-treated and untreated cells (Fig. S6). This suggests that GlcNAc moieties may be involved in an independent mechanism of attachment, which facilitated the residual attachment of *ΔexoY* mutants (Fig. 5A). Altogether, these results indicate that the *exoY* gene is involved in succinoglycan biosynthesis in *P. inhibens* bacteria, and that this polysaccharide is part of the bacterial ECM.

**Figure 6.**
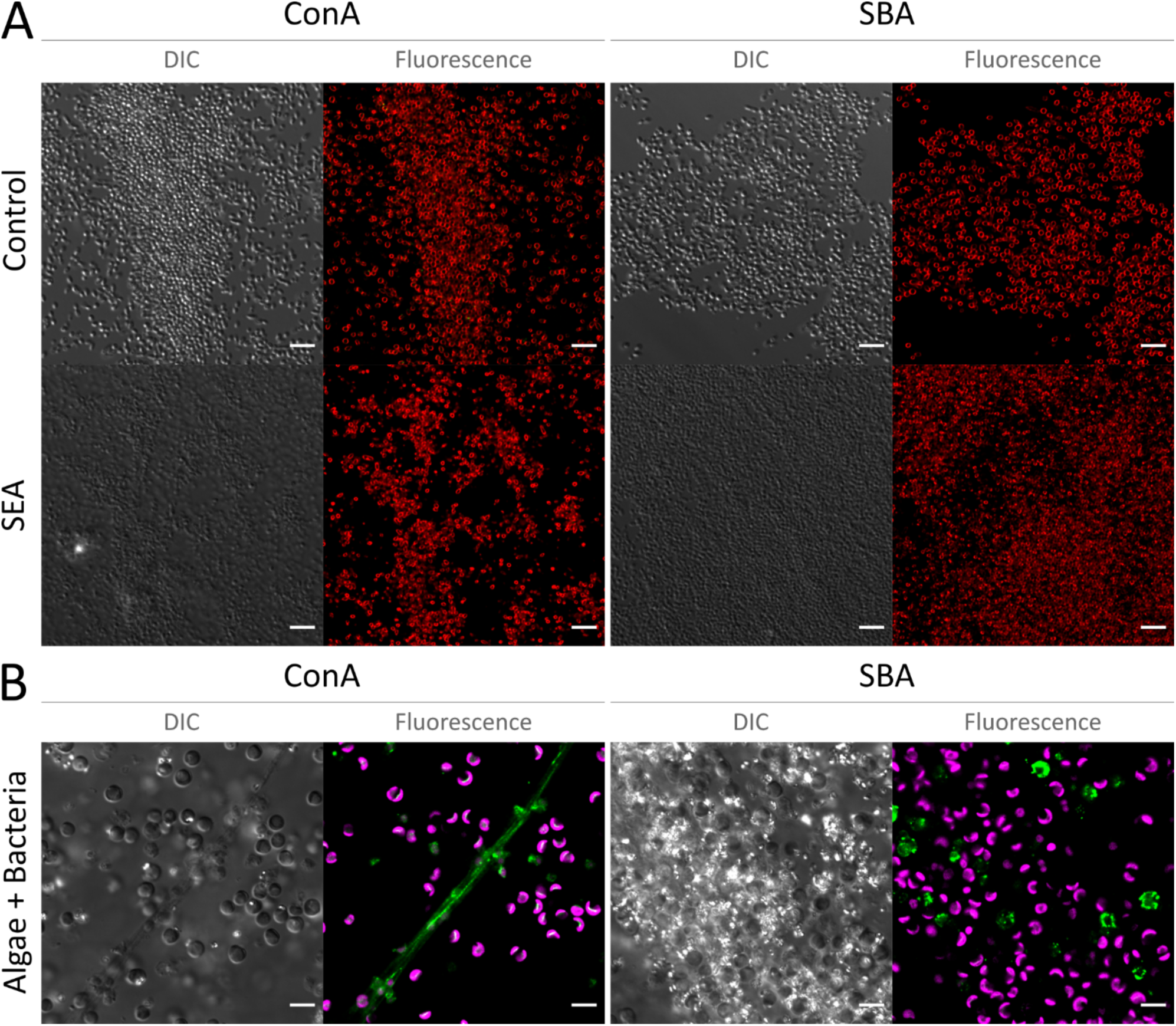
Deletion of the bacterial *exoY* genes results in the absence of succinoglycan-like residues in the joint algal-bacterial ECM. Confocal microscopy was used to visualize the sugar composition of the ECM of different cultures using a panel of FITC-conjugated lectins to detect sugar moieties (green in the fluorescence images). ConA-Concanavalin A that binds glucose residues, and SBA-Soybean Agglutinin that binds galactose residues. **(A)** Bacterial *P. inhibens ΔexoY* cultures were grown to stationary phase on glass slides and treated with SEA (bottom) or were untreated (top). To demarcate bacterial cells the cultures were stained with the membrane dye FM4-64 (red in the fluorescence images). **(B**) *E. huxleyi* -*P. inhibens ΔexoY* co-cultures at late exponential growth. Algal autofluorescence of chlorophyll *a* is shown in pink. Shown are DIC (Differential interference contrast) images for general cell abundance and the fluorescence of the bound FITC-conjugated lectins (green). Scale bar corresponds to 5 µm.

### Co-cultures of algae and *ΔexoY* bacteria produce an ECM with no succinoglycan–like EPS

We now aimed to determine whether the succinoglycan-like EPS in the algal-bacterial ECM is indeed of bacterial origin. Given the perturbed production of EPS in *ΔexoY* bacteria, we generated algal-bacterial co-cultures with mutant *ΔexoY* bacteria and monitored the sugar composition of the joint ECM. Glucose residues (targeted by the ConA lectin), were detected in the ECM (Fig. 6B). Since glucose residues were also detected in algal monocultures (Fig. 4B), this suggests that a polysaccharide with glucose residues is produced and secreted by algae to the extracellular milieu under all tested conditions. However, the polymers with galactose residues that are targeted by the SBA lectin were not detected in the ECM of algal-bacterial co-cultures with *ΔexoY* bacteria. Instead, rare binding in spotted patterns were observed (Fig. 6B), similar to the binding patterns in algal monocultures (Fig. 4B). These results demonstrate that the *exoY* gene product is important for the synthesis of bacterial succinoglycan-like EPS in the joint algal-bacterial ECM.

### Co-cultures of algae and *ΔexoY* bacteria exhibit reduced aggregation

The *ΔexoY* bacteria exhibit perturbed attachment and spatial organization, and co-cultures with these mutant bacteria lack succinoglycan-like residues. To assess the role of bacterial EPS in the aggregation of algal-bacterial co-cultures, we quantified and compared the number of aggregates in co-cultures with WT versus mutant bacteria, or without bacteria. As can be seen in Figure 7A, algal-bacterial co-cultures with *ΔexoY* bacteria produce significantly lower number of aggregates than co-cultures with WT bacteria. Additionally, the cumulative probability of the aggregate sizes showed that the aggregates produced by co-cultures with *ΔexoY* bacteria were significantly smaller than co-cultures with WT bacteria, and similar in size to aggregates produced by algae grown alone (Fig. 7B). These results indicate that bacterial EPS functions as an ECM component that agglomerates algal-bacterial structures.

**Figure 7.**
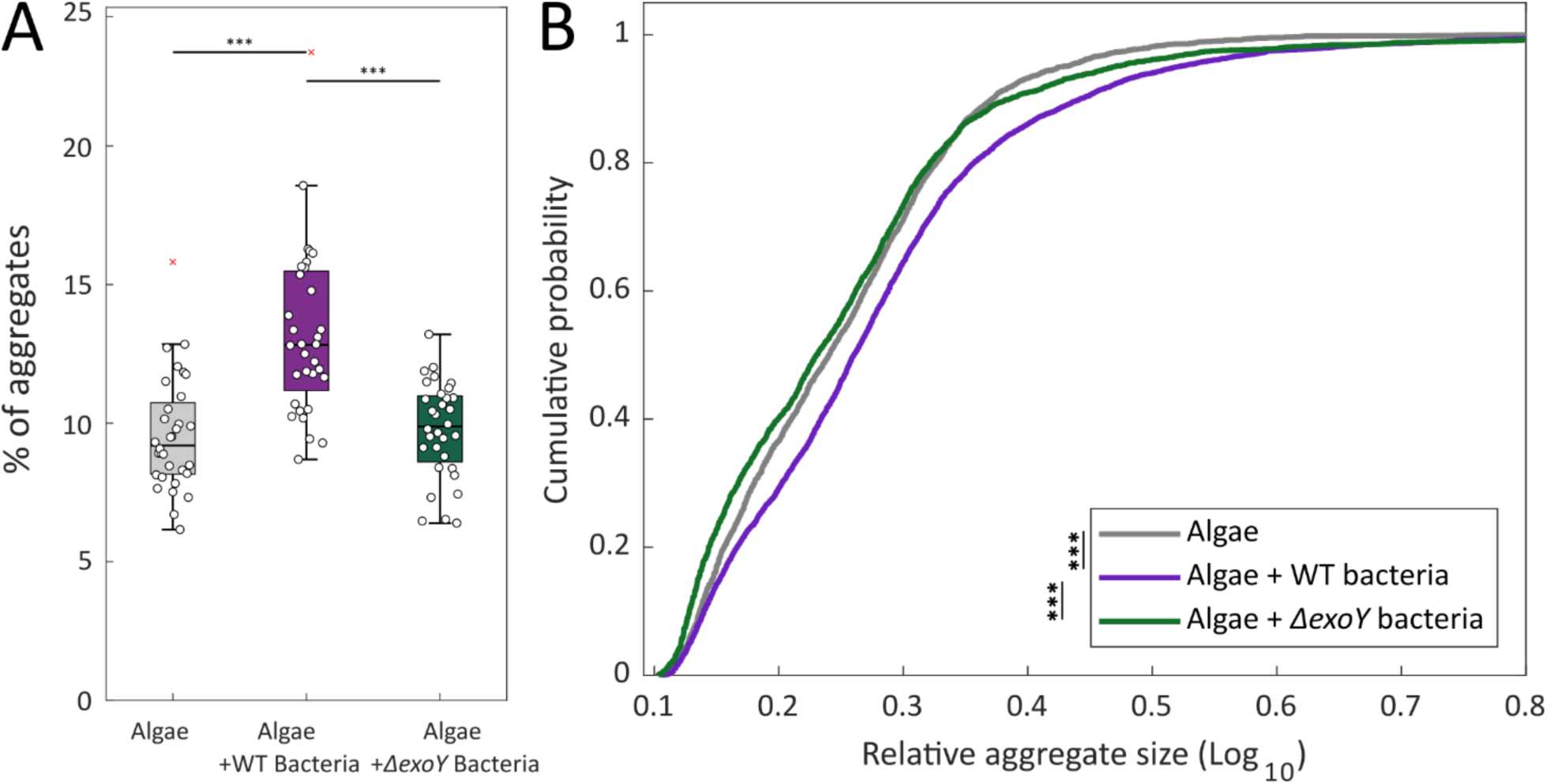
Deletion of the bacterial *exoY_C_* gene reduces aggregation in algal-bacterial co-cultures. Aggregation was measured in cultures of late exponential algal cells without bacteria, algae with WT bacteria and algae with *ΔexoY* bacteria. Aggregation was calculated by imaging algal cells using phase contrast microscopy (see Materials and Methods). Measurements were obtained from n=4 biological repeats, with 8 different images for each biological repeat. **(A)** The percentage of aggregates in the population. Box-plot elements are: center line - median; box limits - upper and lower quartiles; whiskers – min and max values, red x – outlier. Statistical significance was calculated using ANOVA with post-hoc least significant difference (LSD) test. **(B)** Cumulative probability of the relative aggregate sizes (see Materials and Methods). The difference between the cumulative distribution functions was tested using two-sample Kolmogorov-Smirnov tests. Three asterisks denote p-values lower than 0.001.

### Genes involved in wzx/wzy-dependent EPS production are expressed in algal-rich environments

Finally, as a first step to explore the significance of bacterial EPS in algal-bacterial processes in the marine environment, we explored the transcript abundance of genes encoding for proteins involved in bacterial wzx/wzy-dependent EPS production in algal-rich environments. To this end, we analyzed oceanic metatranscriptomics^141^ datasets generated by the TARA Oceans expedition^142,143^. The abundance of transcripts of orthologous groups (OG) of genes involved in wzx/wzy-dependent EPS production was analyzed in the epipelagic layers, as they are known to inhabit phytoplankton^144–147^ and are rich in TEP^148–150^. A positive correlation was observed between transcript abundance of key EPS production genes and chlorophyll *a* concentrations (Fig. 8). Specifically, the functional groups of genes encoding for priming glycotransferases, glycotransferases responsible for repeat unit chain assembly, repeating unit flippases and outer membrane polysaccharide export, showed increasing transcript abundance as chlorophyll *a* concentration increased. Negative correlations, such as in the case of several repeat unit chain glycotransferases could be related to variability in the type of the produced EPS, however these observations require further examination. In summary, it appears that transcripts associated with bacterial EPS production are enriched in algal-rich environments, suggesting a role for bacteria in the formation of joint algal-bacterial ECM and microbial aggregation in these environments.

**Figure 8.**
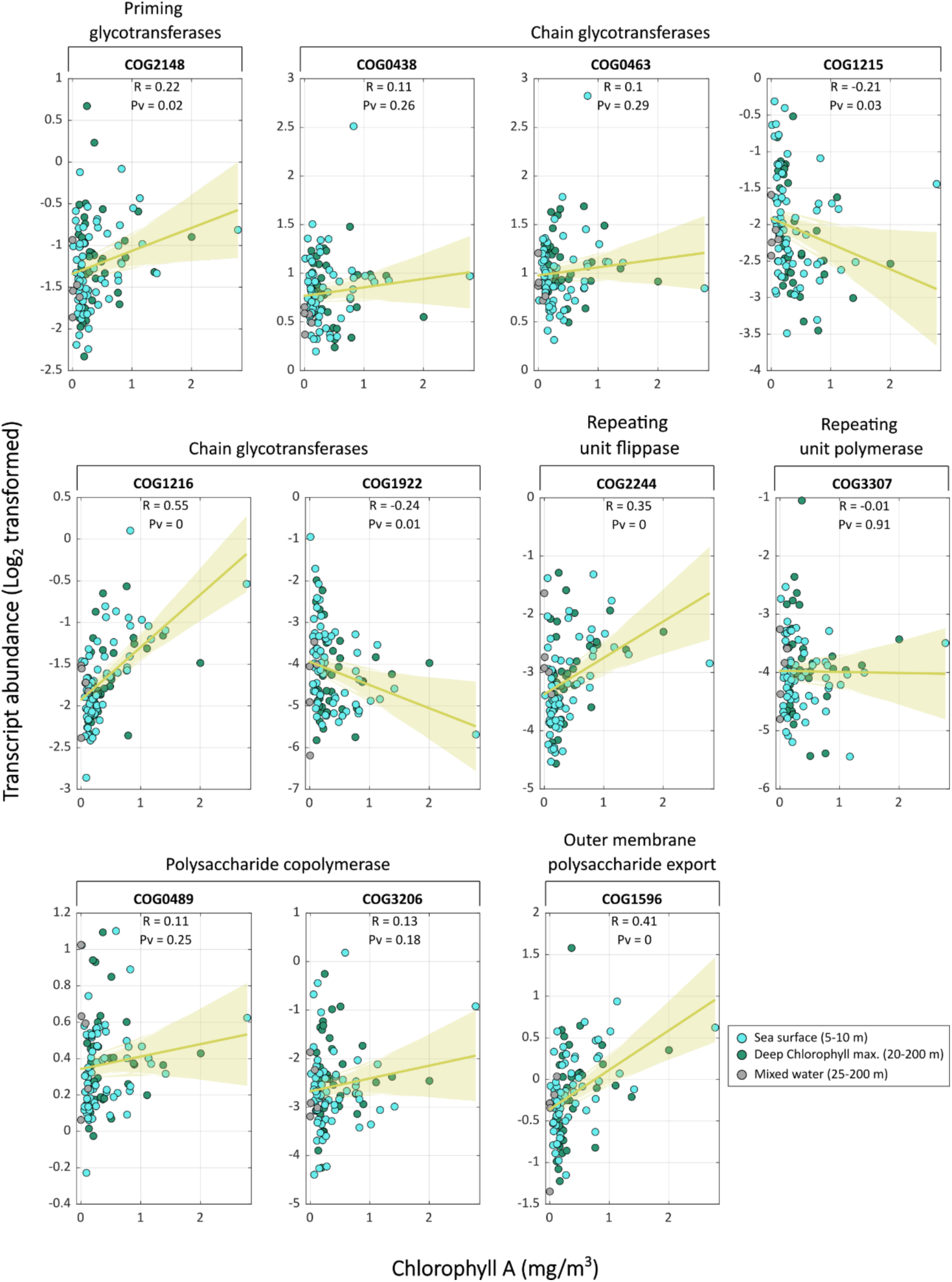
Expression of EPS-related genes in algal-rich layers of the marine environments. Transcript abundances of EggNOG orthologous groups (OG)^216^ of genes encoding for proteins involved in wzx/wzy-dependent EPS biosynthesis in the epipelagic region, collected by the TARA expedition^142,143^. The epipelagic region consists of the sea surface (blue), deep chlorophyll maximum (green) and mixed water layer samples (grey), which originates from the epipelagic region, but cannot be classified to the surface or deep chlorophyll maxima. Transcript abundances (log_2_ transformed) were fit with a linear regression model (yellow line) and a 95% confidence interval (yellow shade) against the chlorophyll *a* concentrations (mg/m^3^) in the sampling point. Shown are R and P-values for each fit. Data adapted from Salazar et al., 2019^141^.

## Discussion

Our data reveals a significant influence of algal exuded molecules on bacterial behavior, facilitating a shift from individual, motile cells to a multicellular lifestyle embedded within an ECM. Previous research has underscored the significance of algal exudates in guiding bacterial chemotaxis towards the phycosphere^77,84,86,151–153^, highlighting the advantages of this function in actively locating and remaining in this microenvironment^77,85^. However the bacterial presence in the phycosphere can be transient and tentative^154,155^, as active mechanisms are required for cells to establish a more enduring connection with a surface^46,156^. While previous studies have shown that impaired chemotaxis affects bacterial attachment to algal cells or algal-produced TEP^33^, the precise impact of algal exudates on the sustained settlement of bacteria in the algal phycosphere remains unclear. In our study, we observed that algal exudates stimulated bacterial attachment to surfaces (Fig. 1) and promoted the formation of complex multidimensional structures (Fig. 2). This was further supported by the concurrent downregulation of motility-related genes and the upregulation of genes potentially involved in attachment or EPS biosynthesis (Fig. 3), demonstrating a transition from free-swimming to a long-lasting, sessile lifestyle^157^. Importantly, these multidimensional structures were found to be held by a bacterially produced ECM (Fig. 4A and S4A), marking the previously unknown contribution of bacteria to the colonization process of algae. Altogether, our findings highlight the multifaceted role of algal exudates, not only in guiding bacterial chemoattraction but also in triggering bacterial processes that promote colonization within the algal microenvironment.

The detection of bacterial EPS associated with the ECM in co-cultures (Fig. 4B) indicates that *P. inhibens* actively produces an ECM within the algal environment. Previous studies have demonstrated the formation of bacterial aggregates on algal ECM surfaces^28,33,158^. Large, extensively colonized TEPs, referred to as ’protobiofilms’, have been shown to play a pivotal role in the early stages of biofilm formation and expedite the establishment of biofilms^28^. In our research, we observed the production of EPS by *E. huxleyi* in both monocultures (Fig. 4B) and co-cultures with either wild type or mutant bacteria lacking EPS production (Fig. 6B). This reinforces the presence of an algal-generated matrix as an available surface for bacterial colonization. Indeed, we observed bacterial cells embedded within TEPs alongside algal cells (Fig. S10), indicating potential attachment of *P. inhibens* to TEPs. The upregulation of bacterial EPS-related genes in co-cultures (Fig. S5B) implies that the formation of bacterial ECM is an integral part of the process of colonizing the algal microenvironment. These findings support the hypothesis that both bacteria and algae contribute to the EPS pool within a shared ECM. In previous studies^28,33,158,159^, bacterial attachment to TEPs was investigated both towards isolated algal TEP and towards a combination of algal TEP and exudates. Our research has illuminated the specific contribution of algal exudates in enhancing bacterial attachment and potentially driving subsequent bacterial EPS production in co-cultures, underscoring the activating influence of algal exudates.

The collaborative formation of an algal-bacterial ECM facilitates the aggregation of both algae and bacteria (Fig. 7). As algal exudates have been shown to influence the recruitment of certain bacteria and establishing symbiotic interactions^86,160^, it is plausible that promoting the formation of a joint ECM can support a close and intimate interaction between algae and bacteria. Numerous studies have illustrated that bacteria can establish close proximity through direct attachment to algal cells^161–163^. Such spatial interactions play a crucial role in mediating nutrient exchange and promoting mutualistic relationships in certain algal species. Research has shown that bacteria directly attached to algal cells assimilate more algal-derived carbon compared to unattached bacteria^64,164^. However, EPS production is energetically demanding^165–167^, and attachment to the algal ECM increases intercellular distances. Therefore, ECM formation in proximity to algal cells likely presents benefits for bacteria. Potential benefits could include opportunities for bacterial cell cooperation^166,167^, including processes such as the extracellular degradation of complex compounds by bacteria^168–171^, as well as the establishment of a shared, enclosed environment with algae. Such a confined microenvironment could provide collective advantages^172,173^ to bacterial populations by enabling the capture of environmental resources, including nutrients and ions^168,169,174^. Furthermore, the creation of localized reservoirs of molecules^175^ may play roles in interspecies and intra-species interactions^59,60,68,176^. The ECM-held algal bacterial arrangement can also benefit algal cells by offering increased resistance to environmental stressors^177^ and protection against predators^178,179^. Furthermore, because bacteria that attach directly to the algal cell have been demonstrated to exhibit pathogenic behavior^59,180–183^, the ECM can serve to create a protective distance, safeguarding algae from bacteria, while still maintaining close proximity for interaction. Thus, both algae and bacteria are likely to benefit from joint algal-bacterial matrix-embedded aggregation.

Uncovering a joint ECM originating from both algae and bacteria invokes important questions about the formation and composition of ECM in marine ecosystems. Our research has revealed that genes related to bacterial EPS are actively transcribed in algae-enriched environments (Fig. 8). However, our understanding of algal-bacterial matrix-building processes in both natural and perturbed environments remains limited. In perturbed environments with excess mucilage where entire ecosystems suffocate due to outbreaks of a thick ECM, the ECM production is predominantly attributed to algae^7–10^, with little discussion of bacterial contributions. Marine mucilage is recognized in encouraging bacterial colonization^184,185^ including pathogenic bacteria^8^. Therefore, it can serve as hot-spots for horizontal gene transfer (HGT) ^186–188^ and the spread of antibiotic resistance genes^189^. With the rising frequency of mucilage events associated with increasing water surface temperatures^8^, gaining a deeper understanding of the algal and bacterial roles played in ECM production is crucial for developing strategies to address future outbreaks.

## Materials and Methods

### Microbial Strains and Growth Conditions

The bacterial strain *Phaeobacter inhibens* DSM 17395 was acquired from the German Collection of Microorganisms and Cell Cultures (DSMZ, Braunschweig, Germany). Bacterial monocultures were cultivated using the following procedure: bacteria were inoculated from a frozen stock (-80°C) onto ½ YTSS agar plates containing 2 g yeast extract, 1.25 g tryptone, and 20 g sea salt per liter (all obtained from Sigma-Aldrich, St. Louis, MO, USA). Plates were incubated at 30 °C for 2 days. Individual colonies were used to initiate bacterial cultures in artificial seawater (ASW) medium. All bacterial monocultures were supplemented with CNPS to support bacterial growth in the absence of algae. CNPS consisted of L1-Si medium (details below) ^59,190^ supplemented with glucose (5.5 mM), Na_2_SO_4_ (33 mM), NH_4_Cl (5 mM), and KH_2_PO_4_ (2 mM), all sourced from Sigma-Aldrich. The cultures were incubated at 30 °C with continuous shaking at 130 rpm for 2 days. Bacterial concentrations were quantified by measuring OD_600_ values using an Ultrospec 2100 pro spectrophotometer (Biochrom, Cambridge, UK) with plastic cuvettes. Cell numbers were estimated based on OD_600_ values and the cultures were diluted to an initial OD_600_ of 0.01 in ASW medium before each experiment. Sampling days of bacterial monocultures are referenced as days following the dilution (day 0): mid exponential (1 day), early stationary (2 days), and stationary (3 days).

The axenic algal strain *Emiliania huxleyi* CCMP3266 (also known as *Gephyrocapsa huxleyi*^191^) was obtained from the National Center for Marine Algae and Microbiota (Bigelow Laboratory for Ocean Sciences, Maine, USA). Algae were cultivated in L1 medium based on the protocol by Guillard and Hargraves^192^. Notably, Na_2_SiO_3_ was omitted as per cultivation recommendations for this algal strain, and the medium was designated as L1-Si. Algal cultures were maintained in standing conditions within a growth room at a temperature of 18 °C under a light/dark cycle of 16/8 h. Light intensity during the illuminated period was maintained at 150 mmoles/m^2^/s. The absence of bacteria in axenic algal cultures was periodically verified through plating on ½ YTSS plates and microscopic observations.

Co-cultures involving *E. huxleyi* and *P. inhibens* were prepared through the following steps: Algal cell concentrations from a late exponential phase culture were enumerated using a CellStream CS-100496 flow cytometer (Merck, Darmstadt, Germany) with 561 nm excitation and 702 nm emission. Each sample involved recording 50,000 events. An inoculum of 10^4^ algal cells was introduced into 30 ml of L1-Si medium and incubated as previously detailed. Following four days of algal growth, *P. inhibens* bacteria (prepared as previously described) were quantified using OD_600_ measurements and cell numbers were adjusted to attain a concentration of 10^2^ cells/ml in 30 ml of algal culture, resulting in 10–100 colony forming units (CFU) per ml. Co-cultures were then incubated under conditions identical to algal cultures. Sampling days of algal monocultures or algal-bacterial co-cultures are referenced as days following the addition of bacteria to co-cultures (day 0): early exponential (4 days), mid exponential (7 days), late exponential (9 days), and early stationary (10 days).

### Monitoring Algal and Bacterial Growth

Algal cell concentrations were quantified using a flow cytometer, as previously described. To ensure accurate measurements, each algal culture was thoroughly mixed to disperse large aggregates. Subsequently, 100 µL of the prepared samples were transferred to round-bottom 96-well plates. A total of 50,000 events were recorded for each sample to determine algal cell counts. Routine bacterial cell concentrations were determined using optical density measurements at 600 nm (OD_600_) in a spectrophotometer, as detailed earlier. For generating growth curves, bacterial cells were appropriately diluted to an initial OD_600_ of 0.01 in the growth 150 µL medium. A layer of 50 µL n-Hexadecane (99%, Alfa Aeser) was added to each well of flat-bottom 96-well plates to prevent evaporation. The bacterial cultures were incubated at 30 °C in an Infinite 200 Pro M Plex plate reader (Tecan Group Ltd., Männedorf, Switzerland). The incubation process involved alternating cycles of 4 minutes of shaking and 25 minutes of incubation, with 300 cycles. Growth was monitored by measuring the absorbance at 600 nm following each shaking step. Growth curves were calculated by subtracting the mean absorbance values of the blanks corresponding to the growth medium. This procedure ensured accurate representation of bacterial growth patterns over time. For co-culture experiments, bacterial concentrations were determined using the colony-forming unit (CFU/mL) method. Bacterial cultures were serially diluted in ASW and plated on ½ YTSS plates. The plates were incubated at 30 °C for 2 days, and subsequently, colonies were enumerated to calculate bacterial concentration in CFU/mL.

### Algal Spent Medium and SEA Treatment

To prepare algal spent medium, axenic algal cultures were grown as previously described for a duration of 7 days. Subsequently, the cultures were filtered using Thermo Scientific™ Nalgene™ Rapid-Flow™ Disposable Filter Units equipped with a polyethersulfone (PES) membrane of 0.2 μm pore size. This filtration step effectively removed algal cells from the culture. The pH of the resulting filtrate was measured and adjusted to pH 8.0 through careful titration using HCl and NaOH solutions. The SEA treatment was formulated to include the following components, each added to the ASW medium, at a final concentration of 1 mM dissolved in MiliQ purified water (unless otherwise stated): dimethylsulfoniopropionate HCl (DMSP, ≥ 96.0%), NaOH, betaine (≥ 99%), disodium succinate, all obtained from Sigma-Aldrich, Merck.

### Attachment Assays

Bacterial cultures were diluted to an optical density of 0.01 at 600 nm (OD_600_) using 150 µL of growth medium. The diluted cultures were dispensed into 96-well flat-bottom plates to initiate the assay. To minimize evaporation during incubation, the plates were securely covered with UV-treated Parafilm. The bacterial cultures were incubated at 30 °C with continuous shaking until the assay endpoint was reached. The attachment assay was carried out following established protocols^79^. After incubation, the plates were subjected to the following steps: The culture medium was carefully aspirated, and the wells were rinsed twice with 1× phosphate-buffered saline (PBS, obtained from Sartorius, Beit HaEmek, Israel). This process aimed to remove non-adherent bacterial cells. The plates were dried by exposing them to a temperature of 55 °C for 20 minutes. The dried plates were submerged in a solution of 0.1% crystal violet. The plates were then incubated at room temperature for 10 minutes to allow the crystal violet to interact with the attached cells. The staining solution was discarded, and the plates were washed twice with PBS to remove excess crystal violet. The plates were left to dry overnight at room temperature to ensure complete fixation of the stained cells. To quantify the attached cells, the remaining crystal violet was extracted by adding 200 µL of 33% acetic acid to each well. The plates were incubated at room temperature for 15 minutes to facilitate extraction. Absorbance measurements were conducted at a wavelength of 595 nm using the TECAN microplate reader. The absorbance values were normalized to a blank consisting of the same growth medium, without bacterial cells, that was subjected to the same protocol.

### Gene Expression Analysis in Pure Bacterial Cultures

To assess the expression of genes associated with motility, attachment, and extracellular polymeric substance (EPS) production, bacterial cultures were grown overnight in ASW medium until they reached an optical density of 0.3 (OD_600_). Subsequently, cultures were subjected to the SEA treatment as detailed earlier or left untreated. Following treatment, cultures were incubated for the specified durations, after which bacterial cells were harvested for RNA extraction. A total of 10^8^ bacterial cells were harvested by centrifugation at 4000 rpm for 10 min. RNA was extracted from the harvested cells using the Isolate II RNA Mini Kit (Meridian Bioscience, London, UK), following the manufacturer’s instructions. Cells were disrupted in RLY buffer containing 1% β-mercaptoethanol using bead beating with 100 µm low-binding silica beads (SPEX, Metuchen, Netherlands) at 30 mHz for 5 min. Approximately 1.4 µg of RNA was treated with 4 µl Turbo DNAse (ThermoFisher) in a 50 µl reaction volume. The RNA samples were cleaned and concentrated using the RNA Clean & Concentrator-5 kit (Zymo Research, Irvine, CA, USA) according to the manufacturer’s instructions.

Equal concentrations of RNA were used for cDNA synthesis employing Superscript IV (ThermoFisher), following the manufacturer’s protocols. qPCR was conducted in 384-well plates using the SensiFAST SYBR Lo-ROX Kit (Meridian Bioscience) on the QuantStudio 5 (384-well plate) qPCR cycler (Applied Biosystems, Foster City, CA, USA). The qPCR program encompassed 40 cycles, adhering to the enzyme requirements. The obtained results were analyzed using the QuantStudio 5 software through a relative standard curve approach. Primer efficiencies were determined by performing qPCR amplification on serially diluted genomic DNA, extracted using the Wizard® Genomic DNA Purification Kit (Promega) as per the manufacturer’s instructions. Only primer pairs with at least 80% efficiency were considered. The expression levels of the target genes were normalized using two housekeeping genes: bacterial housekeeping genes *recA* and *gyrA* (Table 1). No DNA contamination was detected when applying the same program to RNA samples that were not reverse-transcribed. Relative gene expression levels were compared to non-treated samples grown under the same conditions.

**Table 1.**
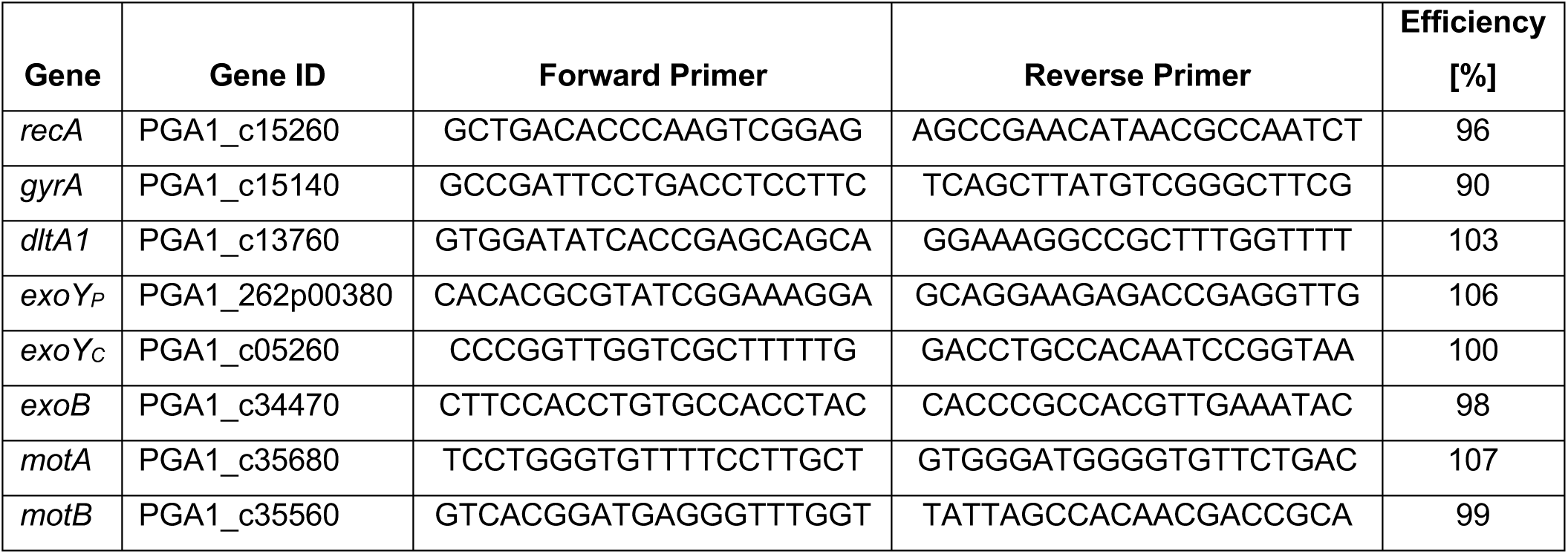
Oligonucleotides used for qPCR of *P. inhibens* genes involved in motility, attachment and EPS biosynthesis.

### Culture Preparation and Staining Procedures for Microscopy

For imaging bacterial biofilms, bacterial cultures were diluted to an optical density of 0.01 at 600 nm (OD_600_) in 12.5 mL of medium. The diluted cultures were grown in Coplin jars containing 7 glass microscopy slides, which had been subjected to dry heat sterilization (170 °C, 3 hours). Parafilm was used to secure the lids of the jars, reducing the potential for evaporation. The jars were then incubated at 30 °C for a duration of 3 days to allow for biofilm formation. Prior to staining, the glass slides were washed twice with phosphate-buffered saline (PBS), and excess liquid was removed by gentle tapping. The staining process involved adding 300 µL of the staining solution (described below) onto the cells. Post-staining, the slides were washed twice with PBS to eliminate any unbound dye. A cover slip was placed over the stained cells, and imaging was carried out promptly.

For imaging algal cultures, co-cultures, or non-surface attached bacteria, the following steps were followed: A 1% agarose solution was prepared in ASW. Small agarose pads were created on glass slides, and the sample of interest was gently placed onto these pads. After a 5-minute air drying period, the agarose pads were subjected to a gentle wash with PBS to remove any excess liquid. The staining solution, as described below, was applied to the samples on the agarose pads. Following staining, the samples were washed twice with PBS to remove excess dye. A cover slip was carefully positioned over the samples on the agarose pads. Subsequently, immediate imaging was performed.

Alcian Blue staining was performed using a staining solution containing 400 mg L^-1^ of Alcian Blue (8GX, Sigma-Aldrich). The solution was prepared in acidified ultrapure water, adjusting the pH to 2.5 with glacial acetic acid. Samples were incubated with the dye in the dark at room temperature for a period of 2 minutes. The staining procedure followed established protocols^193^. Fluorescent dyes were employed at specific concentrations for labeling: Syto9 (Invitrogen™) - 1.7 µM, FM4-64 (Invitrogen™) - 10 μg/mL, FITC-Labeled Lectins (Vector Laboratories) - 20 μg/mL. Samples designated for fluorescent dye labeling were incubated in the dark at room temperature for a period of 30 minutes.

### Microscopy and Imaging Setup

Phase contrast images were captured using a Nikon Eclipse Ni upright microscope fitted with a PLAN 10x Ph1 DL objective lens (Nikon, Tokyo, Japan). Image acquisition was performed utilizing a Nikon DS-Fi3 color camera (Nikon, Tokyo, Japan). For fluorescence and phase contrast imaging, a Nikon Eclipse Ti2-E inverted microscope was employed, featuring a Plan Apo λ 100x Oil Ph3 DM objective lens (Nikon, Tokyo, Japan). Images were acquired using an Andor Zyla VSC-8969 camera controlled by Nis Elements software. Lumencor Spectra X Chroma excitation filters with wavelengths of 470/24 and 575/25 were utilized. FITC-WGA was visualized with an ET519/26 m filter. Laser scanning confocal images were obtained using a Nikon Eclipse Ti2-E A1R HD25 inverted microscope. The system was equipped with PLAN Apo10x λS OFN25 DIC N1 and SR Plan Apo IR AC 60xWI objective lenses (Nikon, Tokyo, Japan). Image acquisition was performed using a Nikon A1 LFOV camera, controlled through Nis Elements software. The scanner selection was Galvano, and the detector selection was DU4 with GaAsP CH2/3 configuration. A first dichroic mirror with wavelengths of 405/488/561/640 was employed. For fluorescence visualization, a 525/50 filter cube was used to capture the fluorescence signal of Syto9 and FITC-labeled lectins. FM4-64 and chlorophyll *a* autofluorescence were captured using 593/46 filter cubes. Consistent image processing protocols were applied across all compared image sets to ensure uniformity.

### Quantification of Biofilm Thickness

Biofilm thickness was quantified using confocal imaging to visualize Syto9-stained bacteria adhered to glass slides. Z-stack confocal imaging was performed with a step size of 0.5 µm. Initial imaging was conducted at an x10 magnification to identify the attachment zone at the liquid-air interface. Subsequently, images were captured at ×60 magnification from seven randomly selected points for detailed analysis. The BiofilmQ tool^194^ was employed for image analysis. The Syto9-stained images were subjected to segmentation using the Otsu thresholding method. Segmentation utilized a cube with a side length of 20 voxels (equivalent to 5.75 µm). From the segmented images, two key biofilm thickness metrics were calculated: (1) Biofilm Mean Thickness: The average thickness of the biofilm was calculated based on the segmented images. (2) Surface Local Thickness: This metric quantified the local thickness of the biofilm. To represent the multidimensional thickness data, particularly surface local thickness, ParaView^195^, a visualization tool, was utilized. ParaView facilitated the creation of visual representations that effectively highlighted variations in biofilm thickness across the imaged area.

### Microscopy and Image Analysis for Aggregate Size Distribution

Samples were cultured as previously described. To initiate imaging, 10 µL of each sample was loaded onto a hemocytometer. For every sample, eight outer corner squares (1 mm²) were imaged using phase-contrast microscopy at a magnification of ×10. To capture the complete dimensions of each square, two images were acquired and subsequently manually stitched and cropped using Inkscape^196^. This approach ensured uniformly sized images for analysis. FIJI ImageJ^197^ software was employed to calibrate the captured images. For segmentation of cells, the Trainable WEKA segmentation plugin^198^ was utilized. The plugin was trained using a dataset comprising 15 randomly cropped images from diverse samples. Binary segmentation images were generated, which served as the basis for further analysis. Particle analysis was conducted using FIJI ImageJ. Particles with sizes < 1 μm² or located on the image border were excluded from analysis. The particle size distribution data, log10 transformed, exhibited three distinct parts in the curve. This included a concave curve representing particle sizes smaller than individual cells, a flat curve reflecting the size distribution of single cells, and a convex curve representing the size distribution of aggregates. To extract the size distribution of single cells, linear fittings were applied to the three curve segments. Two significant change points were identified, characterizing the transition points between these segments. Sizes exceeding 3 median absolute deviations (MAD) between these points were classified as aggregates, while sizes less than 3 MAD were categorized as non-cell particles and discarded. Aggregate sizes were normalized by dividing them by the mean of the size distributions of single cells for each respective image.

### Cation Exchange Resin (CER) Extraction of EPS

The EPS extraction procedure using cation exchange resin (CER) was adapted from established protocols^199^. Cultures were grown in 120 mL of the designated medium, as described previously. For the determination of dry weight (DW), a 5 mL aliquot of the sample was subjected to centrifugation at 3200 ×g and 4°C for 30 minutes. The resulting pellet was re-suspended in 1 mL of 1× PBS, transferred to pre-weighed Eppendorf tubes, and then centrifuged again under identical conditions. Following removal of the supernatant, the pellets were dried overnight at 103°C in an oven. The dried pellets were weighed allowing to estimate DW per culture volume for subsequent CER extraction. For the CER extraction, an aliquot of 100 mL of the sample, estimated to contain approximately 10 mg of DW, was centrifuged at 4000 ×g and 18°C for 15 minutes. The pellet was re-suspended in 1 mL of 1× PBS, and the same centrifugation conditions were repeated to collect cell aggregates. Subsequently, the pellet was suspended in 1 mL of 1× PBS along with 500 mg of CER beads (Amberlite® IR120 Na^+^ form, Sigma-Aldrich), which corresponded to a ratio of 50 grams of cationic beads per gram of DW biomass. This suspension was incubated for 90 minutes on a rotary disc shaker at 150 rotations per minute and 4°C. After the incubation, the suspension was centrifuged (4000 ×g, 15 minutes) to isolate the initial supernatant, and the pellet was re-suspended in 1 mL of 1× PBS. Following another round of centrifugation under the same conditions, the two suspensions were combined. The resultant mixture was subjected to dialysis for two days using MilliQ purified water and SnakeSkin™ Dialysis tubing (3,500 MWCO, Thermo Scientific). The samples were subsequently stored at –80°C and subjected to freeze-drying (Christ LyoCube, Gamma 2-16 LSCplus) for a period of 3 days to concentrate the samples. For carbohydrate quantification, approximately 1 mg of the freeze-dried sample was re-suspended in 0.5 mL of 1× PBS. Total carbohydrate content was determined using the Anthrone method, adapted for microplate reading^200,201^. In triplicates, 50 μL of the sample was mixed with 150 μL of 0.1% Anthrone reagent (ASC, Sigma-Aldrich) dissolved in 96% sulfuric acid (ACS, Sigma-Aldrich). This mixture was incubated in a 96-well microplate at 60°C for 30 minutes, followed by cooling to room temperature for 10 minutes. Polysaccharides were quantified at 620 nm using a TECAN microplate reader. Carbohydrate levels were determined based on a calibration curve derived from glucose standards (ranging from 0 to 100 mg/mL). The amount of carbohydrate per mg of sample DW was calculated using the following equation:

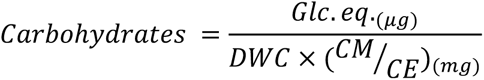

Here, Glc. eq. represents the weight of glucose corresponding to the OD_620_ measurement of the sample, DWC is the dry weight per milliliter of sample, CM is the concentration of the re-suspended sample used for measurements, and CE is the dry weight of extracted EPS per milliliter of sample.

### Functional Annotation of Genes and Phylogenetic Analysis

To elucidate the potential functions of genes surrounding the *exoY* genes in *P. inhibens* (PGA1_c05320 on the chromosome and PGA1_262p00380 on the 262kb plasmid), functional annotation was carried out using EggNOG-mapper v2^114,115^. This mapping process facilitated the assignment of these genes to specific orthologous groups, providing valuable insights into their potential roles. For protein-coding gene families, the protein sequences of the *P. inhibens exoY* genes were aligned using the PANTHER HMM model sequence alignment tool^123^. This approach aimed to identify aligning gene families and to gain information about potential related functions. A dataset of protein sequences was compiled from the UniProt database^202^, focusing on sequences associated with the KEGG orthologous groups (KO) corresponding to the genes present in the producing bacteria. To ensure specificity, only sequences from the Proteobacteria group were retained for subsequent analysis. To focus on priming glycotransferases, the MAFFT algorithm (version 7.520) ^203^ was employed for multiple sequence alignment (MSA), utilizing the FFT-NS-2 strategy. The alignment results underwent visual inspection to identify any misalignments or gaps. Notably, no manual editing or refinement was applied, preserving the integrity of the alignment. Aligned protein sequences were utilized to construct a phylogenetic tree, portraying the evolutionary relationships among these sequences. The PhyML software^204^ (version 3.0) was employed for this purpose, utilizing the maximum likelihood method. To enhance accuracy, a Q.pfam +R+F substitution model was chosen, facilitated by Smart Model Selection for PhyML (SMS)^205^. Visualization of the constructed phylogenetic tree was accomplished using the ATGC PRESTO (a Phylogenetic tREe viSualisaTiOn) tool. Subsequent refinement of the tree visualization was conducted manually using Inkscape^196^, optimizing clarity and presentation.

### Transcript Abundance Analysis in Co-Cultures

Transcript abundance data for EPS-related genes in *P. inhibens* during co-culturing with *E. huxleyi* were extracted from pre-existing transcriptome data^66^. To enable a meaningful comparison of absolute transcript abundances among the genes within the *P. inhibens* EPS modules, we utilized transcripts per million (TPM) normalization method. These values were generated from bacterial feature counts. This approach not only accounts for variations in gene length but also effectively normalizes the data based on sequencing depth. In this study, we focused on evaluating the expression levels of the identified genes within the EPS modules of *P. inhibens*. The temporal expression patterns at mid-exponential phases were compared specifically in two conditions: *E. huxleyi - P. inhibens* co-cultures were analyzed to assess the impact of the interaction between the two organisms. *P. inhibens* monocultures grown on glucose were also analyzed to establish a baseline for comparison.

### Genetic Manipulation of *P. inhibens* Bacteria

The deletion mutant *ΔexoY* (strain ES174) was generated through genetic exchange with the gentamycin resistance gene using double homologous recombination^176^. PCR amplifications and restriction-free cloning^206^ were carried out using Phusion High Fidelity DNA polymerase (Thermo Fisher Scientific, Waltham, MA, USA). PCR products were subsequently purified using the NucleoSpin Gel and PCR Clean-up kit (MACHEREY-NAGEL, Düren, Germany). Plasmids were purified using the QIAprep Spin Miniprep Kit (QIAGEN, Hilden, Germany). First, 3 fragments were PCR amplified with approximately 1,000 bp regions upstream and downstream of the target locus. Primers 1 and 2 were used for amplifying the upstream fragment, while primers 3 and 4 were used for the downstream fragment (Table 2).

**Table 2.**
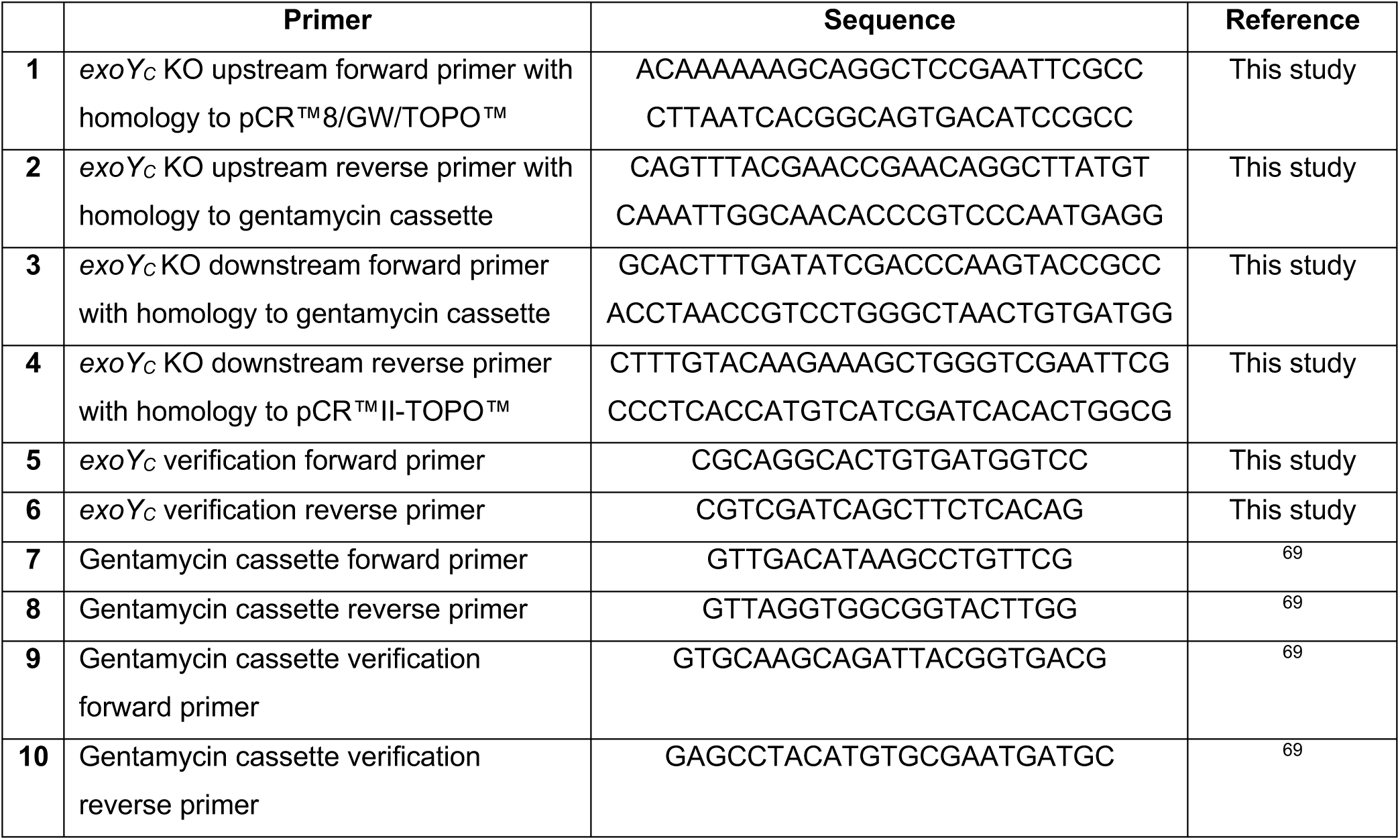
Oligonucleotides used to generate *exoY_C_* KO mutant.

Additionally, a gentamycin resistance gene was amplified from the plasmid pBBR1MCS-5 using primers 7 and 8. The generated PCR products were assembled and cloned into the pCR™8/GW/TOPO™ vector (Invitrogen, Thermo Fisher Scientific, Waltham, MA, USA) using a restriction-free cloning strategy. This resulted in the creation of a knockout (KO) plasmid named pVL1. For transformation, 10 µg of the KO plasmid were mixed with 300 µl of competent *P. inhibens* cells prepared according to established protocols^207^. Competent cells were subjected to electroporation using a pulse of 2.5 kV (MicroPulser, Bio-Rad Laboratories, Hercules, CA, USA), followed by recovery in 2 mL of ½ YTSS medium for 12 hours at 30 °C. Transformed cells were then plated on selective ½ YTSS medium agar plates containing 30 µg/ml of gentamycin. Transformants were screened using single colony PCR with primers 5-6 and 9-10. To confirm the desired mutation, individual cell clones were subjected to Sanger sequencing for verification.

### Transcript Abundance of EPS-Related Genes in Marine Environments

Transcript abundances of EPS genes were functionally annotated in the current study and documented by Salazar et al. 2019^141^, as part of the TARA Oceans consortium study. Briefly, prokaryote-enriched RNA samples were collected from various ocean layers, and paired-end sequencing was conducted using the HiSeq2000 system (Illumina). Sequencing reads from each RNA sample underwent quality filtering and trimming. These reads were then aligned to the functionally annotated Ocean Microbial Reference Gene Catalog v2. Read counts were normalized based on gene length, summarized according to EggNOG gene families (aligned to the EggNOG version 3 database^208^), and divided by the transcript abundance of a constitutively expressed marker gene. DESeq2 was employed for variance stabilization. The resulting transcript abundances, represented as log_2_-transformed values indicating relative transcript numbers per cell, have been made publicly accessible (https://www.ocean-microbiome.org/). For our analysis, we focused on epipelagic ocean layers, including the surface, mix, and deep chlorophyll maximum layers. In conjunction with the transcript abundance data, we incorporated chlorophyll *a* concentration measurements obtained from the respective samples (https://doi.org/10.5281/zenodo.3473199). Using Matlab^209^, we employed the default linear model (fitlm) to fit the transcript abundances within the context of the available data. This process enabled us to derive insights into the relationship between transcript abundances of EPS genes and chlorophyll *a* concentrations in different ocean layers.

## Acknowledgments

We appreciate the technical guidance of Dr. Shifra Ben-Dor, and are thankful for the help of Dr. Ron Rotkopf with statistical analysis (Life Sciences Core Facilities, Weizmann Institute of Science, Israel). We thank Professor Edo Bar-Zeev (Ben-Gurion University, Israel) for sharing his expertise in confocal microscopy imaging. We are thankful to all members of the Segev lab for insightful discussions and comments. This study was supported by funds received from the Weizmann SAERI program, the European Research Council (ERC StG 101075514), the Israeli Science Foundation (ISF 947/18), the Minerva Foundation with funding from the Federal German Ministry for Education and Research, and the de Botton Center for Marine Science, granted to E.S.

## Authors Contributions

V.L. and E.S. designed the study. V.L. and O.S. performed and analyzed experiments. J.R. assisted in mentoring the study. V.L. and E.S. wrote the manuscript. All authors discussed the results and contributed to the final manuscript.

## Supplemental Data

**Figure S1.**
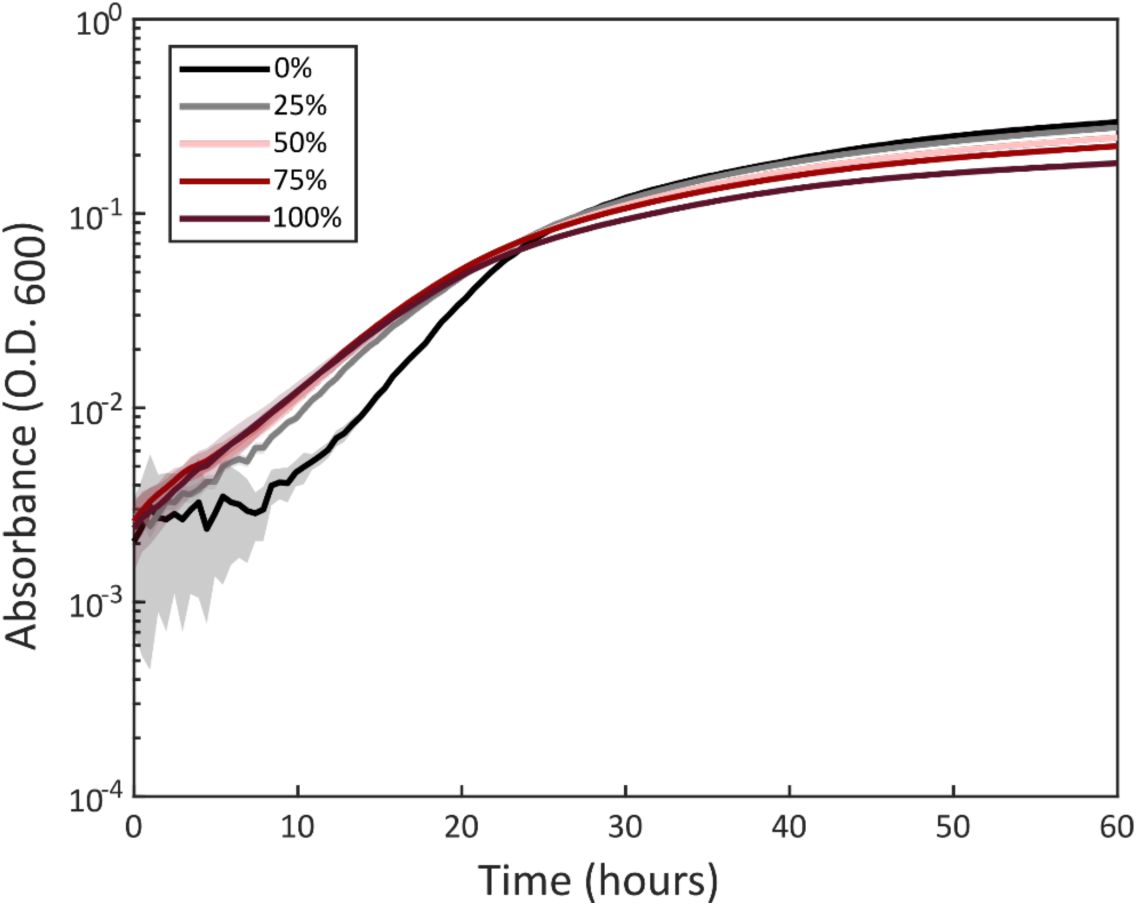
Addition of *E. huxleyi* algal spent medium does not increase the maximum yield of *P. inhibens* bacteria. Growth of *P. inhibens* bacteria was measured by O.D._600_ in 0-100% (v/v) concentrations of spent medium extracted from exponentially growing *E. huxleyi* algal cultures, mixed with ASW. Lines represent the mean of n=6 biological repeats with shaded error bars. All concentrations of spent medium induced lag-phase shortening compared to ASW (0%), as previously described^69^. However, the maximum O.D. was not increased upon treatment with algal spent medium and was even slightly lower at higher spent medium concentrations.

**Figure S2.**
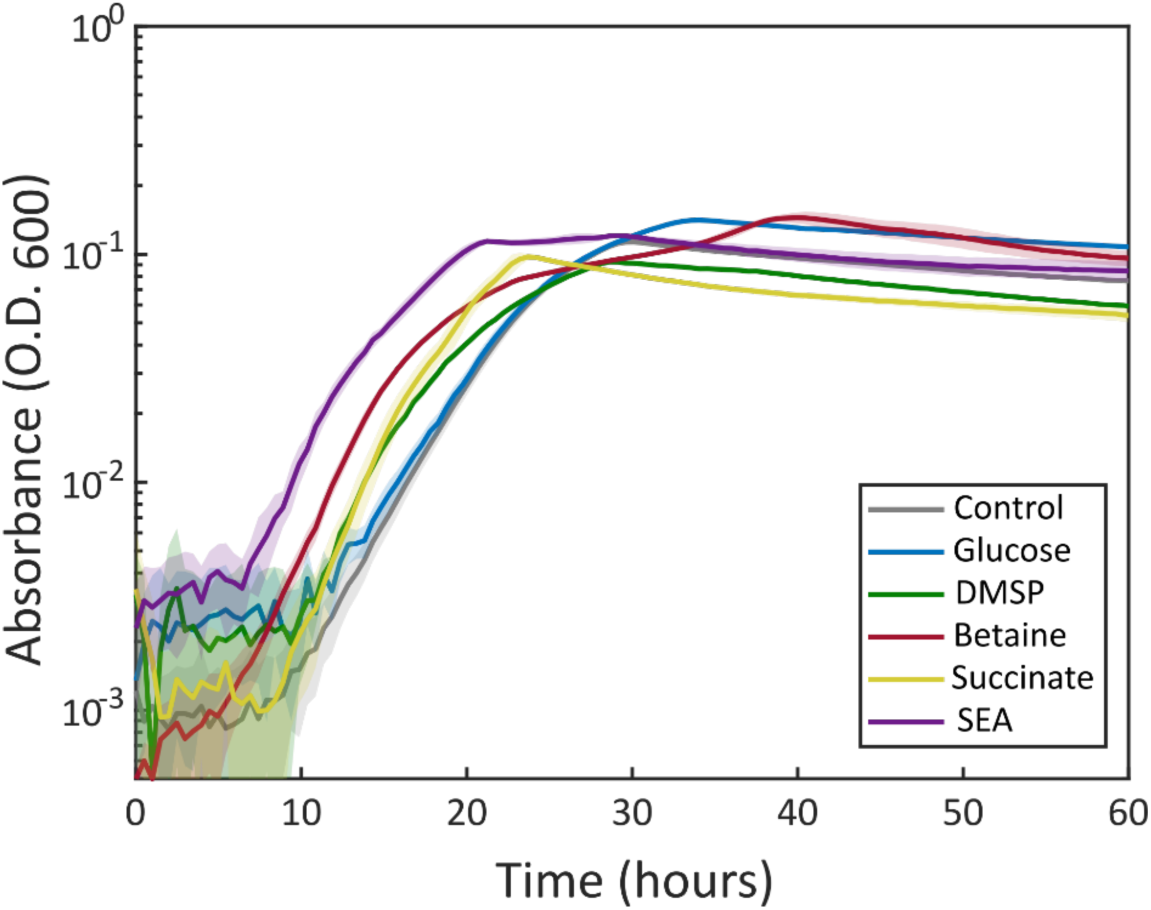
Addition of known *E. huxleyi* algal exudates does not increase the maximum yield of *P. inhibens* bacteria. *P. inhibens* bacteria were supplemented with 1 mM of DMSP, betaine, succinate, or a mixture of 1 mM of each compound (the SEA mix). As controls, bacteria were grown without any supplements or with 0.6 mM glucose, to allow comparison with the amount of carbon in succinate. Growth was measured by O.D._600_. Lines represent the mean of n = 6 biological repeats with shaded error bars. DMSP, betaine, succinate and SEA expedited the bacterial lag-phase compared to the control, as previously described ^69^. However, the maximum O.D. of bacteria supplemented with SEA, DMSP and succinate was equal or slightly lower than the control.

**Figure S3.**
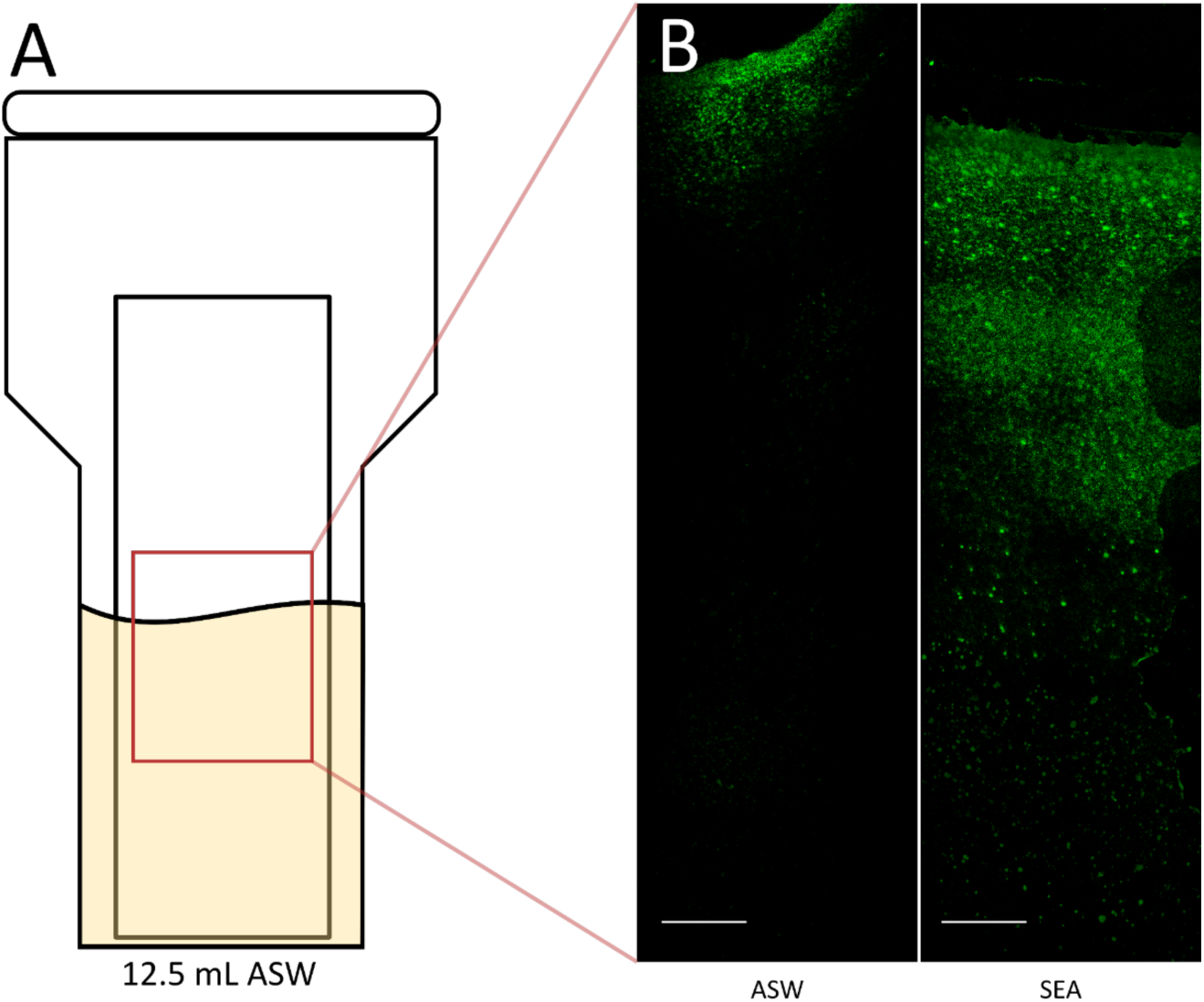
SEA treatment increased *P. inhibens* attachment on glass slides. (A) *P. inhibens* bacteria were grown in Coplin jars and glass slides were inserted to the jars to serve as a surface for attachment. Bacteria were supplemented with SEA (1 mM each compound), or untreated, and grown to stationary phase. (B) Representative stitched confocal images of the glass slides (×10 magnification). For visualization and quantification, each slide was stained with Syto9 (green). As can be seen, bacterial structures mainly formed at the liquid-air interface. Scale bars represent 1000 µm.

**Figure S4.**
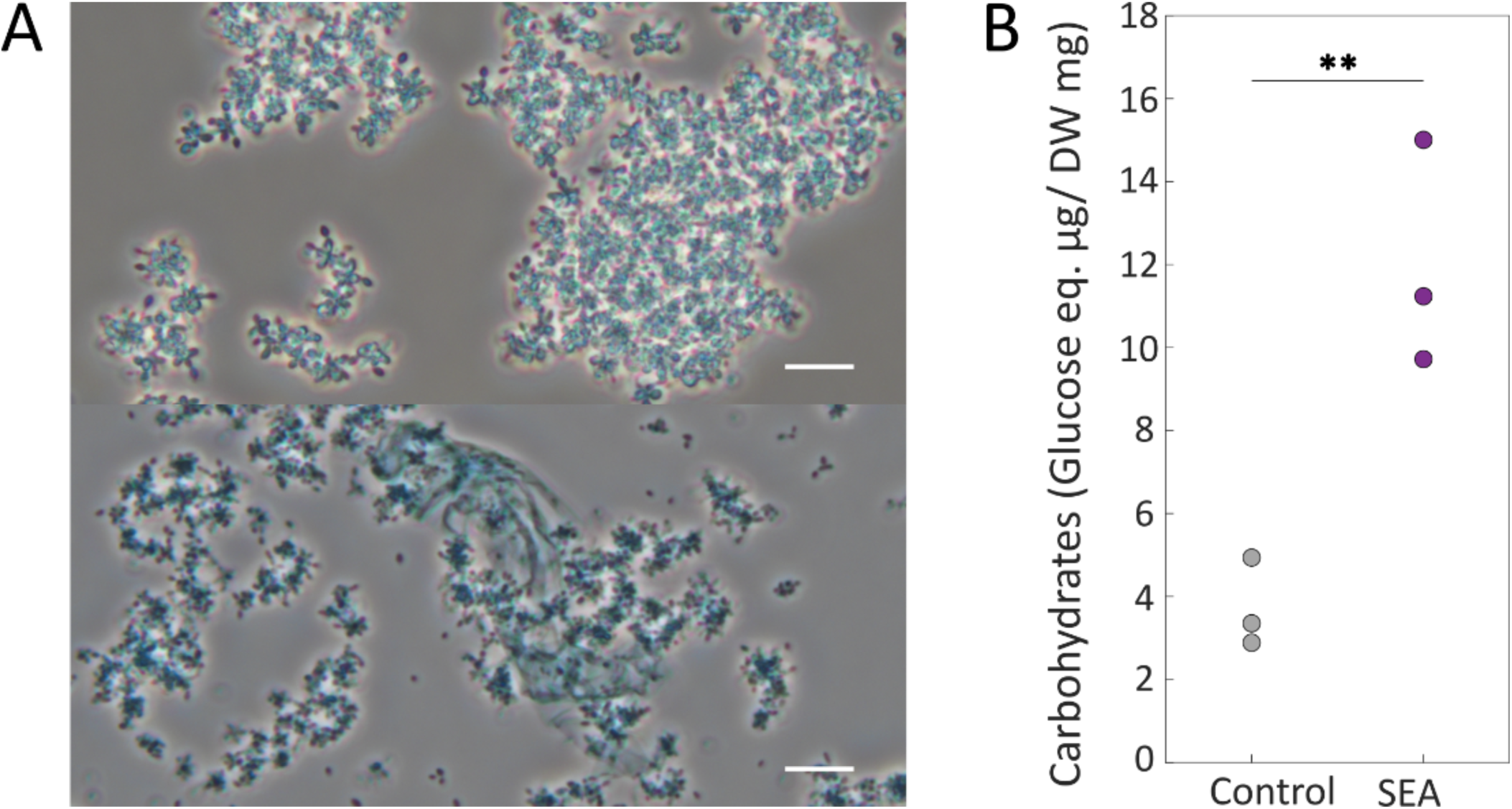
Algal exudates promote bacterial extracellular matrix (ECM) production. **(A)** Bacterial cultures at early stationary phase were stained with Alcian Blue to detect extracellular acidic polysaccharides. SEA-treated bacteria exhibited extracellular sheet-like structures that were stained (bottom, marked by a white arrowheads) while untreated cells did not exhibit similar structures (top). Scale bar represent 20 µm. **(B)** Quantification of extracellular carbohydrates extracted from SEA treated (purple) or untreated (gray) bacteria at stationary phase. For each treatment, carbohydrates were extracted from n=3 samples. Amount of carbohydrates is represented by the equivalent weight of glucose, normalized to the dry weight (DW) of the sample. Statistical significance was calculated using two sample t-test, two asterisks denote p-value < 0.01.

**Figure S5.**
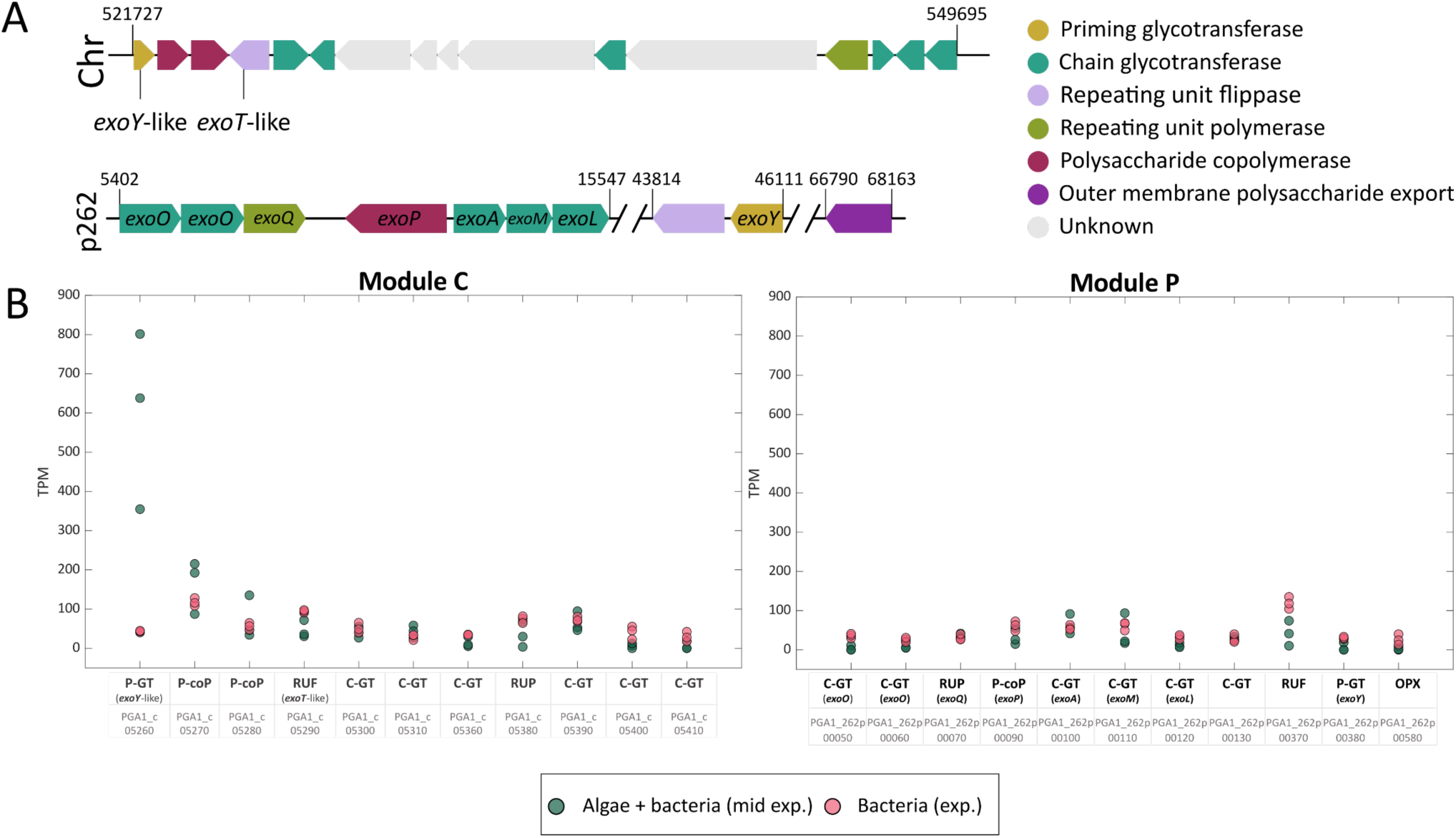
*P. inhibens* EPS module expression in co-cultures. **(A)** *P. inhibens* genes encoding for proteins potentially involved in succinoglycan-like EPS production are clustered on the chromosome (module C) and the p262 kb native plasmid (module P). **(B)** Expression of the identified EPS-related was analyzed in *P. inhibens* bacteria at exponential growth, grown in co-culture with *E. huxleyi* algae (earlt exponential, green) or in monocultures (mid exponential, red). For Each gene, the number of transcripts per million bacterial cells (TPM) was calculated from n=3 biological replicates. Shown is the functional annotation for each gene (P-GT – priming glycotransferases, C-GT – chain glycotransferases, RUF – repeating unit flippase, RUP – repeating unit polymerase, P-coP – polysaccharide co-polymerase, OPX – outer membrane polysaccharide export) and below the gene entry.

**Figure S6.**
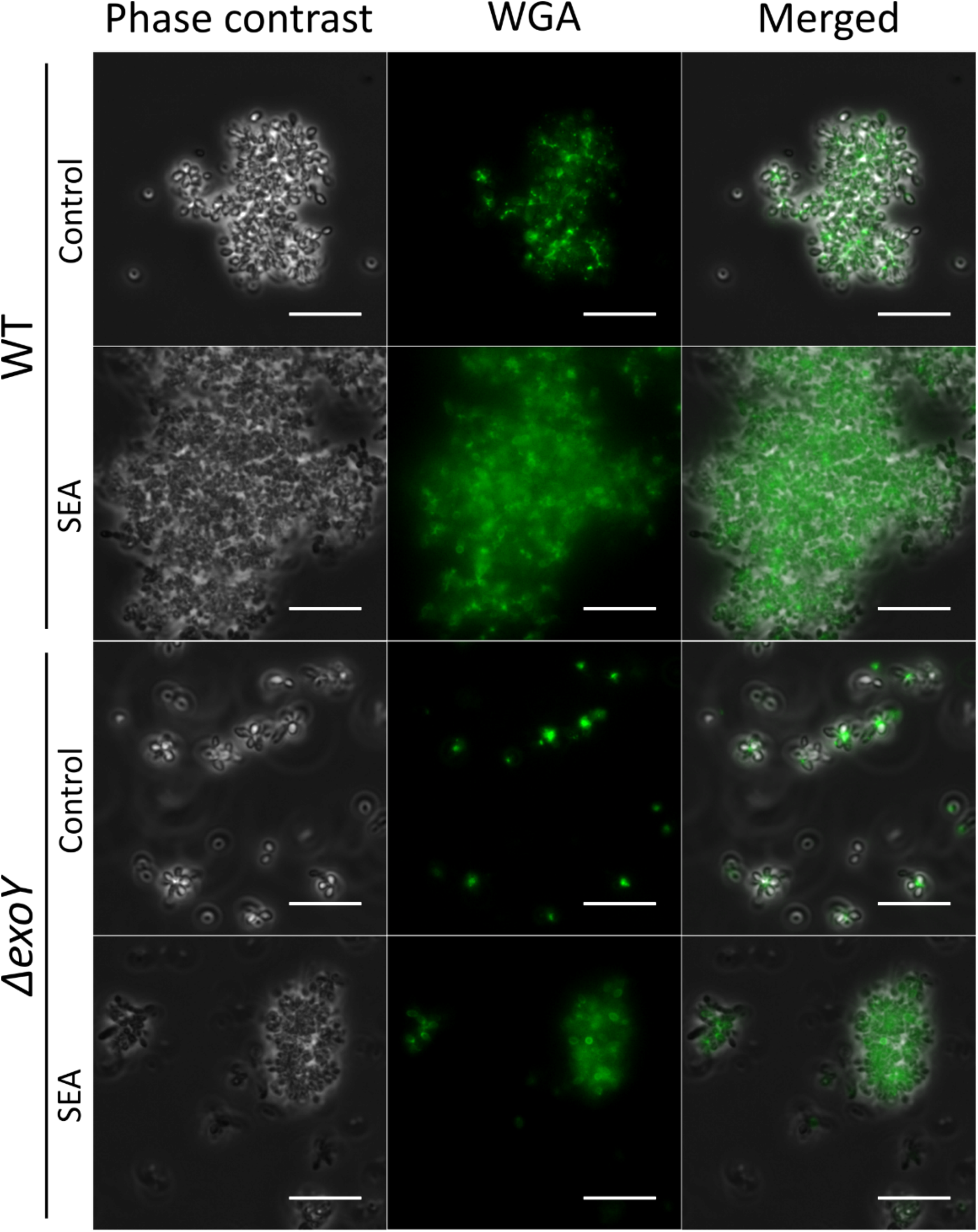
Known algal exudates affect the WGA lectin binding of *P. inhibens*. The polar polysaccharide of *P. inhibens* WT or *ΔexoY* (grown to stationary phase) was stained by FITC-conjugated WGA lectin (green). Both WT and *ΔexoY* showed polar staining in the center of rosettes when cells were untreated. Upon SEA treatment (1 mM of each molecule) WGA lectin exhibited binding to whole cell, in both WT and *ΔexoY* bacteria. Scale bars correspond to 10 µm.

**Figure S7.**
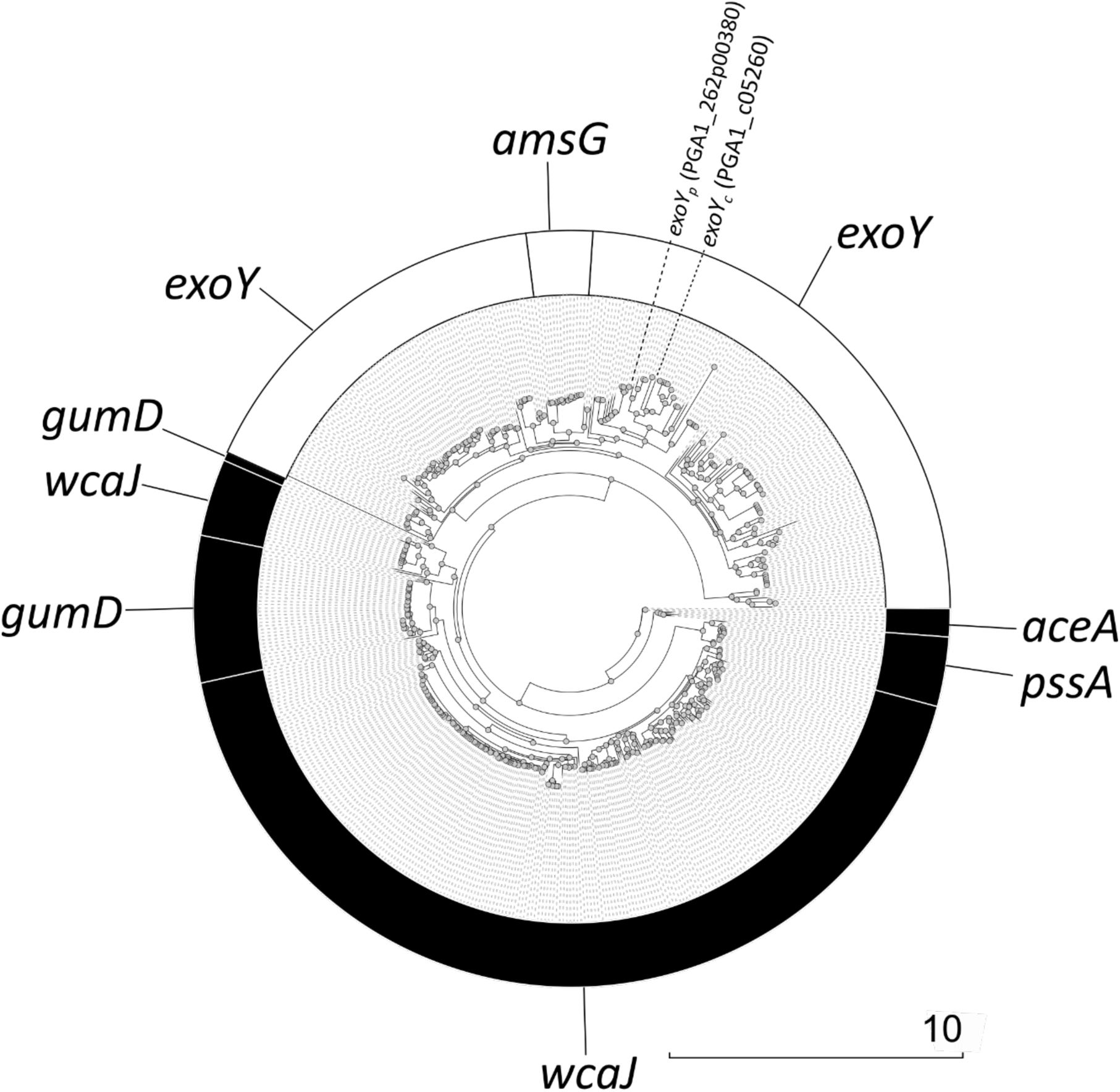
Phylogenetic tree of priming glycosyltransferases of different wzx/wzy-dependent EPS. The protein sequences encoded by the genes of the priming glycotransferases of characterized EPS (Table S2) from *Proteobacteria* were used to perform a multiple sequence alignment. A phylogenetic tree was reconstructed to study the clustering of UDP-glucose (black) and UDP-galactose (white) transferases. The two *P. inhibens exoY* genes are marked inside the UDP-galactose transferases clusters.

**Figure S8.**
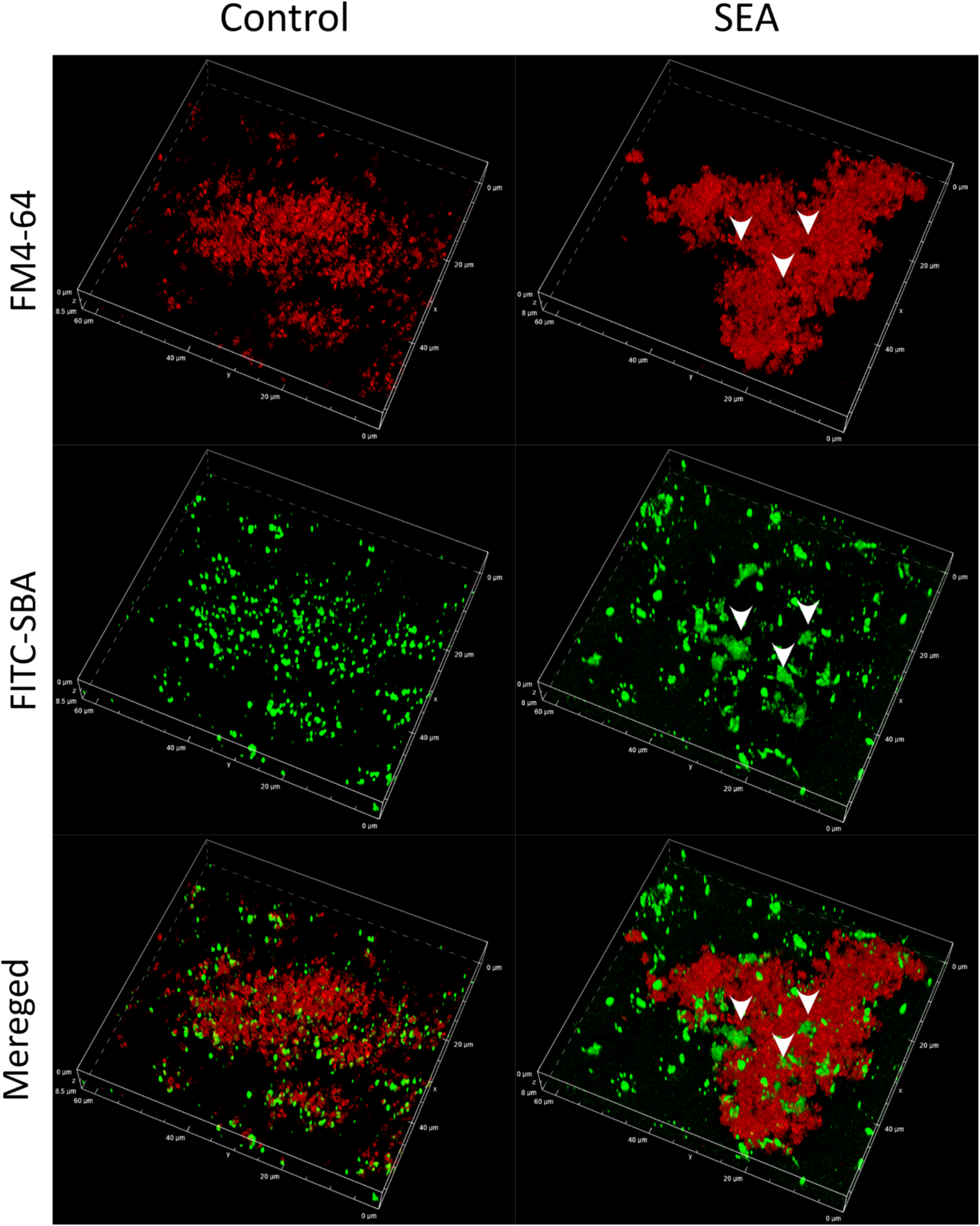
Known algal exudates affect the SBA lectin binding of *P. inhibens*. Volume configuration of the confocal microscopy image shown in Fig. 4A (bottom panels). Bacterial *P. inhibens* monocultures were grown to stationary phase on glass slides and treated with SEA (right) or were untreated (left). Shown are the fluorescence of the bound FITC-conjugated Soybean Agglutinin (SBA, green) that binds galactose residues and the membrane dye FM4-64 (red) that demarcates bacterial cells. White arrowheads show examples of FITC-SBA binding to regions that appear to fill intercellular spaces (see cell distribution according to the membrane stain, FM4-64).

**Figure S9.**
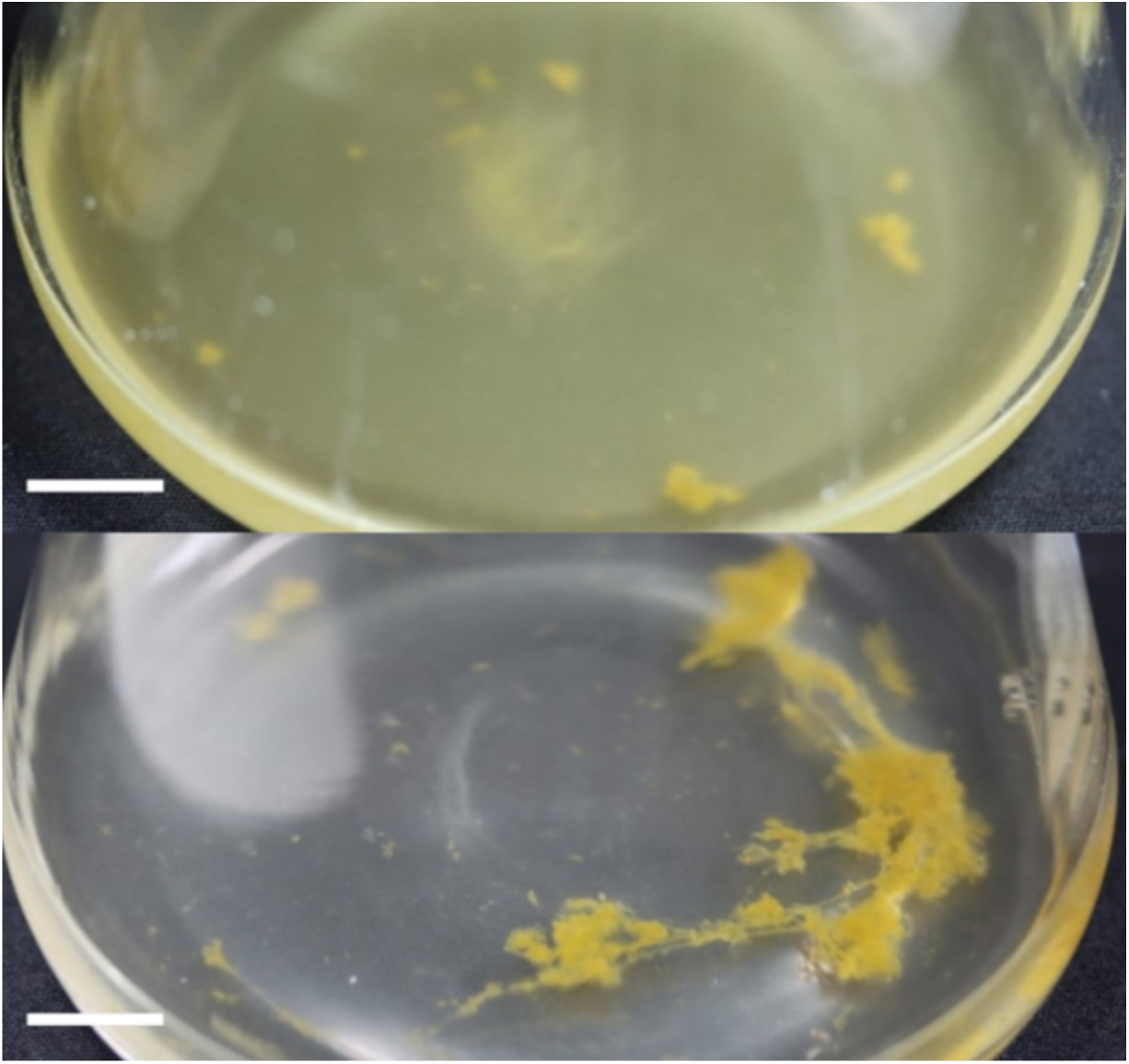
*P. inhibens- E. huxleyi* co-cultures show higher aggregation compared to *E. huxleyi* monocultures. *E. huxleyi* monocultures grown to early stationary phase show dispersed cultures with few aggregates (top), while *P. inhibens-E. huxleyi* co-cultures of the same age show visibly higher aggregation. Scale bar represents 10 mm.

**Figure S10.**
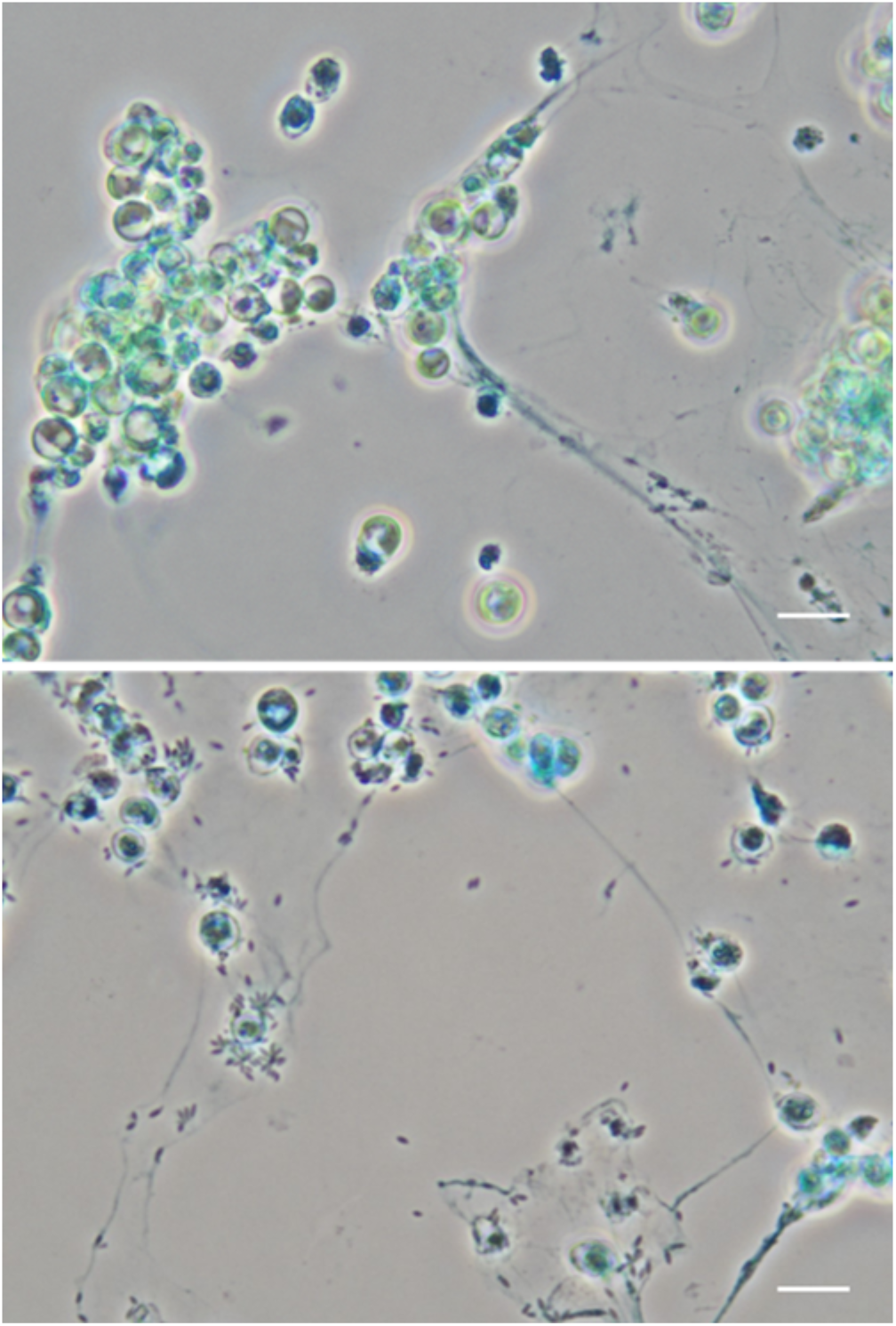
Algae and bacteria produce an extracellular matrix (ECM). Algal and algal - bacterial cultures were stained with Alcian Blue to stain extracellular acidic polysaccharides. Stained threads are seen both in cultures of early stationary phase *E. huxleyi* monocultures (top) and *E. huxleyi*-*P. inhibens* co-cultures (bottom). In co-cultures, *P. inhibens* bacteria are seen in close proximities to the threads. Scale bar represent 20 µm.

**Figure S11.**
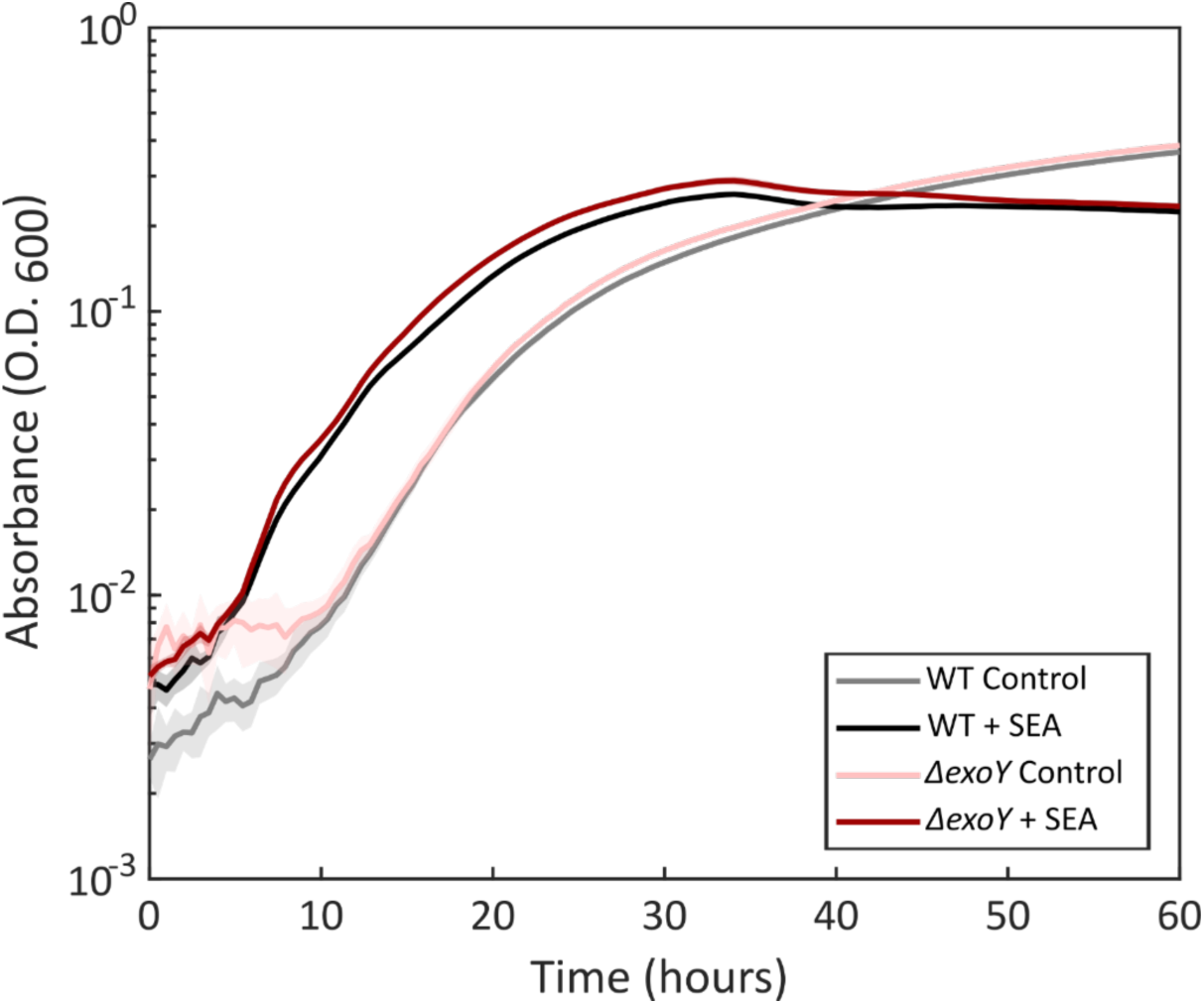
*ΔexoY* bacteria exhibit growth dynamics similar to WT bacteria. *P. inhibens* WT or *ΔexoY* bacteria were supplemented with SEA (1 mM each molecule) or untreated. Growth was measured by O.D._600_. Lines represent the mean of n = 4 biological repeats with shaded error bars. *ΔexoY* mutant showed similar growth to WT both when supplemented with SEA or untreated. both *ΔexoY* and WT bacteria showed shorter lag-phases in response to the SEA treatment, as previously described^69^. Maximum O.D. reached by *ΔexoY* was similar to WT both for SEA-treated and untreated cultures.

**Table S1.**
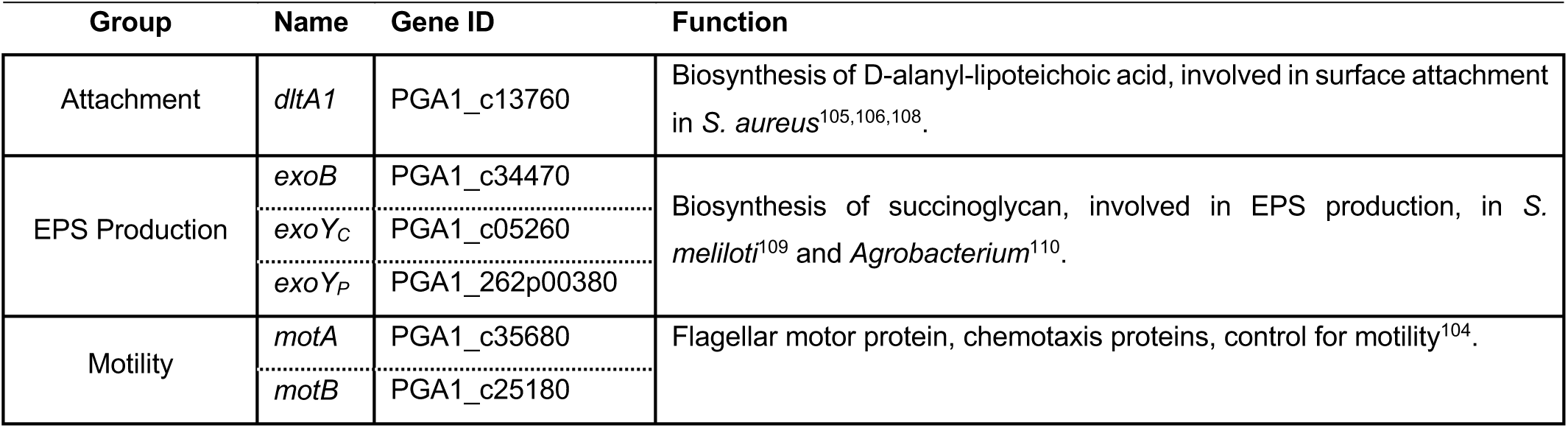
Genes related to key processes during the transition of *P. inhibens* bacteria from motile to sessile states. Genes potentially involved in motility, attachment and EPS production were selected based on homologous pathways in other characterized bacteria.

| Group | Name | Gene ID | Function |
| --- | --- | --- | --- |
| Attachment | *dltA1* | PGA1_c13760 | Biosynthesis of D-alanyl-lipoteichoic acid, involved in surface attachment in *S. aureus*<sup>105,106,108</sup>. |
| EPS Production | *exoB* | PGA1_c34470 | Biosynthesis of succinoglycan, involved in EPS production, in *S. meliloti*<sup>109</sup> and *Agrobacterium*<sup>110</sup>. |
|  | *exoY<sub>C</sub>* | PGA1_c05260 |  |
|  | *exoY<sub>P</sub>* | PGA1_262p00380 |  |
| Motility | *motA* | PGA1_c35680 | Flagellar motor protein, chemotaxis proteins, control for motility<sup>104</sup>. |
|  | *motB* | PGA1_c25180 |  |

**Table S2.**
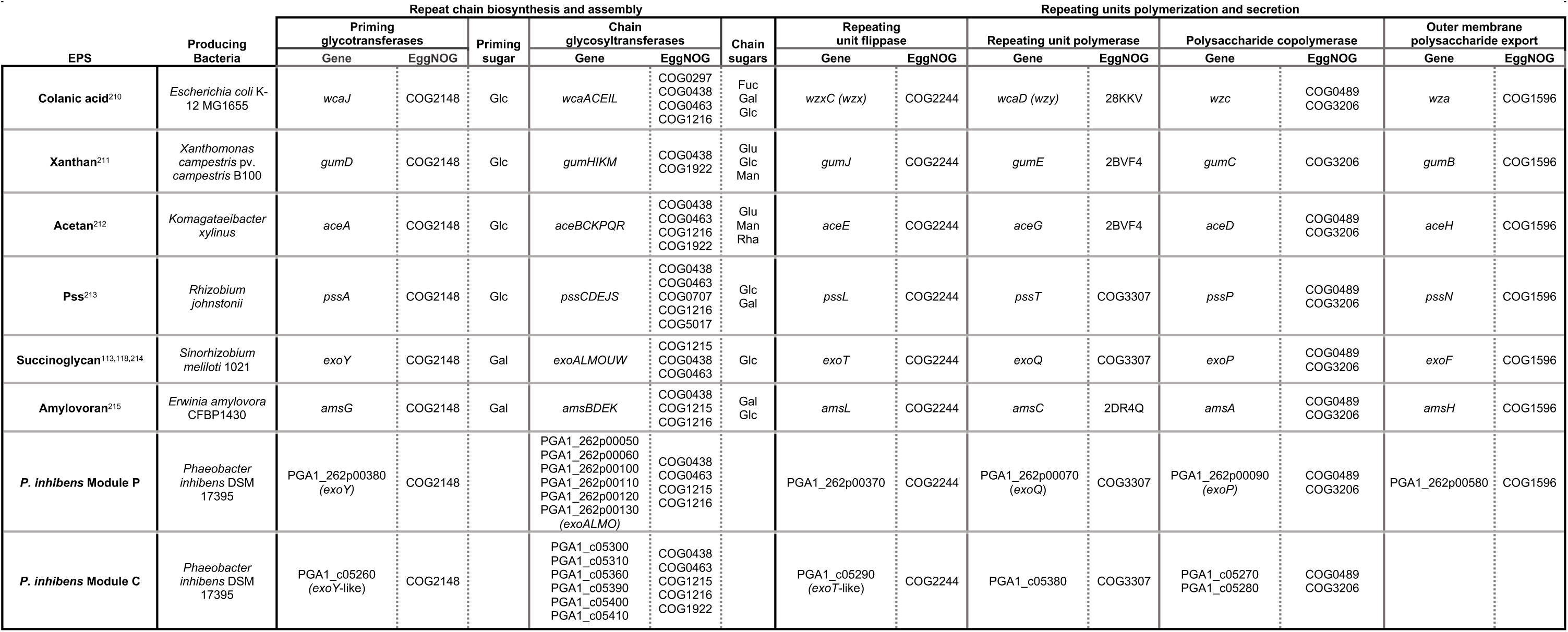
Genetic modules of wzx/wzy-dependent exopolysaccharides. Genes of characterized wzx/wzy-dependent EPS biosynthesis modules including colanic acid, xanthan, acetan, *Rhizobial* Pss, succinoglygan and amylovoran, in key producing bacteria and the corresponding genes in the two *P. inhibens* modules P and C (found on the native 262kb plasmid and on the chromosome, respectively). Shown are the names and functional annotation of genes involved in addition of the priming sugar to the lipid carrier, repeat sugar chain biosynthesis, polymerization of the repeating units and transport. Functional annotations were generated using EggNOG-mapper v2^114,115^, and the identified clusters of orthologous groups (COG) number is specified.

